# Heteroplasmy and tandem repeats reveal adaptation to elevation in the New World Jays (Aves: Corvidae)

**DOI:** 10.1101/2023.12.05.570157

**Authors:** Flavia Termignoni-Garcia, Katia Bougiouri, Scott V. Edwards

## Abstract

Advances in high-throughput sequencing (HTS) and bioinformatic tools have enabled the quick and cost-efficient assembly of complete mitochondrial genomes (mitogenomes) in non-model organisms. Consequently, new evidence of heteroplasmy, recombination and paternal leakage in mitogenomes has increased. In this study, we utilized HTS data from whole-genome sequencing to assemble the first complete mitogenomes of nine species of New World Jays (NWJs), covering all genera. We further investigated the evolution of heteroplasmy, tandem repeats (TRs) and signatures of natural selection. Our results showed a molecular shift in the adaptation to low elevation in the NWJs. Among the species studied, we found 10 heteroplasmic sites either containing TRs in the same site or 1 to 300 nucleotides adjacent; one species-specific TR in a transfer RNA (tRNA-P) potentially associated with low elevation; one phylogenetic branch with evidence of episodic positive selection also associated with low elevation; and 5 codon sites with strong support for positive selection. We referred to the heteroplasmy-TR interaction and its possible role with regulation, recombination and paternal leakage in the mitogenomes. Finally, phylogenetic relationships were in agreement with previous studies and we discussed how selective pressure on genes from the oxidative phosphorylation pathway (OXPHOS) may benefit species from low-elevation habitats. Although these findings in the NWJs require further investigation, this study offers promising insights about the evolution of mitogenomes in birds.

## INTRODUCTION

The mitochondrial genome (mitogenome) is a valuable molecular tool to study biodiversity and evolution. Mitochondrial DNA (mtDNA) has been used for phylogenetic studies, reconstructing biogeographic histories, in population genetics (Avise et al., 1987; Searle, 2000; Boore et al., 2005; Avise, 2009; Paijmans et al., 2013), as a tool in animal conservation (du Toit et al., 2017; Tikochinski et al., 2020), to study its pivotal role in adaptation (Zhou et al., 2014; Morales et al., 2015; Lamb et al., 2018; Ramos et al., 2018; Sebastian et al 2020; Zhao et al 2022) and mitochondrial fitness to understand health and diseases (Song et al., 2014; Sebastián et al., 2017; Stier et al., 2019; Wang et al., 2020). Advantages of using mtDNA include its high mutation and low recombination rate, high number of copies, relatively ease with which it can be isolated, and predominantly maternal inheritance (Kvist, 2003; Abbott et al., 2005; He et al., 2013; Alexander et al., 2015; Halley et al., 2015), as well at its vital role in energy production through the oxidative phosphorylation (OXPHOS) system (Gershoni et al., 2009).

Although mtDNA has been extensively used in many fields of biology, some aspects of mitogenomes still need to be better understood (reviewed by Nicholls & Minczuk, 2014). Its pivotal role in adaptation has been demonstrated, with evidence of climate propelling mitochondrial selection found in Australian songbirds (Morales et al., 2015; Lamb et al., 2018), penguins (Ramos et al., 2018) and Galliforms (Zhou et al., 2014), indicating its potential in other systems. Another aspect of mitogenomes that require further comprehension is the extend of recombination. For example, we know that mitogenomes are minimally recombinant (Hoarau et al., 2002; Tatarenkov & Avise, 2007), though mitogenomes assembled from high-throughput sequencing (HTS) and long-read sequencing (LRS) have revealed that recombination can occur in long blocks, the degree to which it occurs is still debated (Sammler et al., 2011; Wang et al., 2015; Thapana et al., 2022; Ye et al., 2022).

Additionally, the discovery of paternal leakage of mtDNA in birds and other taxa (Kvist, 2003; Abbott et al., 2005; He et al., 2013; Alexander et al., 2015; Halley et al., 2015; Gandolfi et al., 2017; Luo et al., 2018) suggests that mitogenomes may not be primarily maternally inherited as previously thought. Likewise, although heteroplasmy - the sequence and structural variation of mitogenomes within an individual - was thought to be rare in the PCR era, with HTS technology it seems to be prevalent in many taxa (van den Ameele et al., 2020; reviewed in Parakatselaki & Ladoukakis, 2021; Pereira et al., 2021); suggesting its role as a mechanism that provides genetic diversity in natural populations (Huang et al., 2019; Tikochinski et al., 2020; Pereira et al., 2021). Finally, tandem repeats (TRs) in mitogenomes, in conjunction with other processes such as regulation and recombination, likely drive the extreme variability of the mitochondrial control regions (Mundy et al., 1996a; Casane & Guéride, 2002; Hoarau et al., 2002; Ray, 2003; Mundy & Helbig, 2004; Mjelle et al., 2008; Omote et al., 2013; He et al., 2013; Padhi, 2014). However, further research is needed to understand the role of TRs in mitogenome evolution and adaptation.

Genome skimming from HTS for assembling mitogenomes is cost-efficient and quick (Hahn et al., 2013; Gan et al., 2014; Dierckxsens et al., 2016). An additional advantage of using HTS for mitogenome assembly, is that the process captures the total mtDNA from whole-genome sequencing and overcomes the hurdles of older methods, such as long-range PCR. Older methods can produce inaccuracies due to the amplification of nuclear mitochondrial DNA segments (NUMTs), the variation of sequences at primer bonding sites and changes in gene structure (Mueller & Boore, 2005; Cameron, 2014; Parakatselaki et al., 2022). On the other hand, mitogenomes still present challenges that only LRS can overcome, such as accurately capturing gene arrangements, long TRs and long duplications (Konrad et al., 2017; Torres et al., 2019; Zardoya, 2020; Montaña-Lozano et al., 2022).

The higher frequency of mtDNA in animal tissues relative to nuclear DNA (Shadel & Clayton, 1997) also facilitates mitogenome skimming from HTS, making it possible to obtain high read coverage even from low concentrations or from degraded DNA, such as from museum and ancient DNA samples (Guschanski et al., 2013; Paijmans et al., 2013; Anmarkrud & Lifjeld, 2017). In birds, the high mitochondrial density in flight muscles (Kovacs & Meyers, 2000; Butler, 2016) makes them an excellent source of mitogenomes in avian genomic libraries.

In the present study, we assemble complete mitogenomes from HTS data from nine species of New World Jays (NWJs), with at least one representative of each genus. The NWJs, a monophyletic group from the Corvidae family which lack publicly available complete mitogenomes, comprise seven genera (*Aphelocoma, Calocitta, Cyanocitta, Cyanocorax, Cyanolyca, Gymnorhinus* and *Psilorhinus*) (de los Monteros & Cracraft, 1997; Ericson et al., 2005) and around 36 species (Madge, 1994). They constitute an interesting study system because they exhibit a great diversity of reproductive behaviors. They also occur in widely divergent habitats, ranging from warm tropical forests to temperate climates and high altitudes. We utilize the NWJ mitogenomes to investigate levels of heteroplasmy, TRs and evidence for natural selection. We make use of a phylogenetic framework to better understand the evolution of NWJ mitogenomes. Overall, we hope our findings advance the discussion of heteroplasmy-TR interaction, regulation, adaptation and recombination in non-model avian mitogenomes.

## MATERIALS AND METHODS

### Samples

We included in this study a total of 17 samples representing nine species and seven genera of NWJs: the Yucatan jay (*Cyanocorax yucatanicus*) (n=5), the Green jay (*Cyanocorax yncas*) (n=3), the Brown jay (*Psilorhinus morio*) (n=3), the Bushy-crested jay (*Cyanocorax melanocyaneus*) (n=1), the Steller’s jay (*Cyanocitta stelleri*) (n=1), the Unicolored jay (*Aphelocoma unicolor*) (n=1), the Azure-hooded jay (*Cyanolyca cucullata*) (n=1), the Pinyon jay (*Gymnorhinus cyanocephalus*) (n=1) and the Black-throated magpie jay (*Calocitta colliei*) (n=1) (Table S1). We used the publicly available mitogenomes of the Eurasian magpie (*Pica pica*) as the outgroup (NCBI accession number: NC_015200; Liu et al., 2018) and the American crow (*Corvus brachyrhynchos*) (NCBI accession number NC_026461.1; Li et al., 2016) as the bait sequence to retrieve mtDNA reads from HTS.

### DNA isolation, library preparation, and sequencing

We extracted whole genomic DNA from muscle tissue preserved either in ethanol, flash frozen in liquid nitrogen or in *RNALater^®^* RNA Stabilization Reagent (Qiagen, Hilden, Germany) (see Table S1 for details). We used the MagAttract*^®^* HMW DNA kit and the DNeasy Blood and Tissue kit (Qiagen, Hilden, Germany) for DNA isolation. We quantified the purified DNA with a Qubit instrument (Thermo Fischer Scientific) to shear 330 ng of DNA into fragments of ∼ 300 bp with a Covaris sonicator (Covaris Inc.) and, assessed fragment size with Aligent 4200 TapeStation System. We prepared for most of the samples the whole-genome libraries with the PrepXTM ILM 32i DNA Library Protocol and the Apollo Library Prep System, whereas for the Azure-hooded jay, Pinyon jay, and Black-throated magpie jay we used the Kapa HyperPlus kit at ¼ volume. Libraries for the Unicolored jay, Bushy-crested jay and Steller’s jay were sequenced on the Illumina HiSeq (2 x 125 bp paired-end reads) at 50x coverage using one lane. The remaining libraries were sequenced using NovaSeq (2 x 150 bp paired-end reads) at >19x coverage using one lane and at 5x coverage in one lane for the Azure-hooded jay, Pinyon jay and Black-throated magpie jay. All sequencing took place at the Bauer Core Facility at Harvard University. We carried out quality control for all sequenced libraries using FASTQC (Andrews, 2010) and adapters removal with Trimmomatik (Bolger et al., 2014). Most of the samples were initially sequenced for nuclear whole-genome assembly or re-sequencing.

### Mitogenome assembly, circularity inference and annotation

We used three different approaches to assess the accuracy and repeatability of recovering mitogenomes from HTS. In the first approach, we used the program MITObim v1.9 (Hahn et al., 2013), a mitochondrial baiting and iterative mapping method that reconstructs complete mitogenomes from HTS data. This method has been shown to reconstruct high-quality mitogenomes even from relatively low-coverage data (Machado et al., 2016).

Assuming that mtDNA has a random distribution within sequenced libraries, we partitioned the library of every sample sequenced at high coverage into four equal subsets to reduce computational power, time and, to enhance MITObim’s performance. We ran MITObim separately on each of the four subsets of reads per sample (e.g., 01, 02, 03, 04; see Table S2) using the American crow as a bait sequence with the “-quick” option to bypass high memory consumption, and the “-pair” option to use complete read pairs. We compared the four subset assemblies to assess differences and validate the mitogenomes for each sample.

For the second approach we used the AWA software (Machado et al., 2018) on the baited mitogenomes generated by MITObim to infer sequence circularity and complete assembly. This software seeks putative overlapping sequences throughout the mitogenome assembly with the aim of reconstructing a new circular mitogenome sequence with adjacent ends. AWA uses Bowtie2 (Langmead et al., 2009) to map the original reads to the new putative mitogenome. This validates the inferred circularization and provides quality metrics for the assembly, where alignment scores of 3 or lower should be approached with caution (Machado et al., 2018).

For the third approach, we used NOVOPlasty, which assembles organelle genomes; detects heteroplasmy and putative NUMTs; and reconstructs the sequence for each mitochondrial haplotype (Dierckxsens et al., 2016, 2020). NOVOPlasty takes as initial input all the raw reads and sub-sample a fraction of them, depending on the available computation memory. We used a maximum memory of 100 Gb to capture 100% of the input reads during the assemblies. We ran NOVOPlasty using a kmer of 33, and the American crow mitogenome as a bait sequence.

We annotated all mitogenomes with the MITOS WebServer (Bernt et al., 2013) using default parameters and manual inspection. Due to the high variability of the control region (CR), MITOS did not produce specified annotations for this region. Thus, in Geneious v 2019.0.3, we used publicly available CR sequences from species of NWJs, aligned them to our mitogenome and manually annotated a CR for each species. For Unicolored, Steller’s, Black-throated magpie, Brown, Green and Pinyon jays we used the control region of the same species with NCBI accession numbers (respectively): AF218920, AF218924.1, AF218925.1, AF218927.1, AF218930.1, AF218932.1 (Saunders & Edwards, 2000). We used the CR of the closest of the above-mentioned species for the remaining NWJs without a publicly available CR. Finally, we used the assembled mitogenomes produced by NOVOPlasty for downstream analyses, because this assembler provided consistent assemblies and uses all the HTS input reads.

### Alignment and phylogenetic analyses

We performed alignment and manual curation separately for each protein-coding gene (PCG) using MUSCLE v3.6 (Edgar, 2004) and Geneious Prime v 2019.0.3. To maintain frame consistency in alignments, we accounted for the coding nature of sequences by eliminating all normally occurring stop codons, reversed-complemented the ND6 gene and repeated the sequences when necessary for overlapping genes.

We performed phylogenetic analysis on four datasets: one concatenated dataset containing only the 13 PCGs; a second by adding the two ribosomal RNAs (rRNAs); a third by adding the CR; and a fourth consisting of the whole mitogenome. We used the Eurasian magpie mitogenome as the outgroup and estimated sequence divergence (uncorrected p-distances) in MEGA (Tamura et al., 2013) for a rapid test on the feasibility of the assembled mitogenomes.

Prior to phylogenetic analysis, we carried out nucleotide substitution model and selection tests on all four datasets using an optimal partitioning scheme selected by PartitionFinder2 v2.1.1 (Lanfear et al., 2012). We partitioned the 13 PCGs into 39 blocks, with one block for each codon position of each PCG. We grouped the two rRNAs into two separate blocks, the CR sequence in another block and all transfer RNAs (tRNAs) in an additional block. We performed phylogenetic analysis with MrBayes v3.2.6 (Ronquist et al., 2012) in four runs, using four sampling chains, each running for 10^6^ generations, sampling every 100 generations. We discarded the first 35% of trees as burn-in, and a 50% majority rule consensus tree was produced from the posterior distribution of trees.

We parsed sequences and alignments to calculate base composition with the toolkit SeqKit (Shen et al., 2016). Base composition can reflect major changes in mitogenomes (Gibson et al., 2005), can be influenced by repeats or non-coding unique sequences that expand the CR (Fauron et al., 1976; Wolstenholme, 1992) and finally influence the mitogenome size. Thus, we also looked for a relationship between base composition and mitogenome length while taking into account phylogenetic relatedness. For this analysis we used Phylogenetic Generalized Least Squares (PGLS) (Symonds & Blomberg, 2014) with the ultrametric tree generated above and assumed the traits evolve under a Brownian motion process (BM) or an Ornstein-Uhlenbeck (OU) model. The R packages we used for these analyses were *ape* (Paradis and Schliep 2019)*, nlme* (Pinheiro 2022) and *phytools* (Revell, 2012; latest version, 1.0-3).

### Heteroplasmy and detection of NUMTs

We detected heteroplasmy and excluded NUMTs with NOVOPlasty (Dierckxsens et al., 2016; 2020). One of the advantages of NOVOPlasty is that it can detect NUMTs during heteroplasmy analysis (Dierckxsens et al., 2020); it detects low-frequency intra-individual polymorphisms and length during a second round of assembly (around the polymorphism) with the raw reads after mitogenome circularization. The double assembly aims to determine whether the haplotype is a NUMT or has mitochondrial origin (true heteroplasmy). We followed the recommendations for organelle coverage in heteroplasmy detection to minimize false positives (Dierckxsens et al., 2020). We used a minor allele frequency (MAF) of 0.01 for organelle average coverage above 500X and, MAF of 0.1 for coverage below 300X. In addition, we conducted multiple analyses to determine levels of heteroplasmy in each sample by gradually increasing the number of reads and memory (Gb) tenfold, starting from 10% and ending with 100% of the total reads.

We compared the positions of the heteroplasmic sites between the NWJ mitogenomes by manually annotating the heteroplasmic sites on each mitogenome with Geneious Prime v 2019.0.3 and later performing a multiple alignment with MUSCLE v3.6 (Edgar, 2004). We used the R packages ggplot2 (Wickham, 2016) and gggenes v. 0.4.1 (Wilkins, 2020) to visualize the number and position of heteroplasmic sites.

### Analysis of tandem repeats

After performing a multiple alignment of whole-mitogenomes, we examined tandem repeats (TRs) on each mitogenome. We utilized Tandem Repeats Finder (trf) v2.02 (Benson, 1999) with a mismatch probability of 75 and a minimum alignment score of 20. We used ggplot2, gggenes, and phytools (Revell, 2012 v.1.0-3) to visualize the number, position, and ancestral reconstruction of TR features, including length, copy number, and quantity. The ancestral reconstruction was conducted using the contMap and fastAnc functions. This process estimates states at internal nodes using maximum likelihoods (ML) and interpolates along edges according to the Felsenstein (1985) equation. We examined the relationship between habitat elevation and TR length for TRs that exhibited in the ancestral reconstruction analyses a specific pattern within clades or convergence. To examine the relationship between TR length and elevation, we took into account any possible dependence among species due to phylogenetic relatedness with PGLS (Symonds & Blomberg, 2014) using the ultrametric tree generated above and analyzed it under both Brownian and Ornstein-Uhlenbeck (OU) models. We used the reported elevation of each specimen as a value for its corresponding subspecies, recognizing that some species undergo speciation under niche divergence (e.g., in the Steller’s jay by Cicero et al., 2022; in the Unicolored jay by McCormack et al., 2008, 2009; in the Yucatan jay by Termignoni-Garcia et al., 2017), while for other species is unknown.

### Analysis of natural selection

We employed MEGA (Tamura et al., 2013) to estimate pairwise and overall synonymous (Ks) and nonsynonymous (Ka) substitutions for each PCG, along with their ratio represented by the omega value (ω = Ka/Ks). A value of ω greater than 1 indicates evidence of positive selection in a gene, while values lower than 1 suggest purifying selection. Even if a gene exhibits ω<1, it may still show signs of positive selection at the codon level. Therefore, we conducted additional analyses to detect selection at codon sites using the programs HyPhy (Kosakovsky et al., 2020) and PAML v4.9h (Yang, 2007).

We used a concatenated dataset containing the PCGs alignment without stop codons (HyPhy rejects alignments with stop codons) and the inferred Bayesian tree as input files for the natural selection analyses. We used one sample per species during analyses to balance the signal across branches. We used two different methods in HyPhy to test for sites under positive or purifying selection: the Fast Unconstrained Bayesian Approximation (FUBAR) (Murrell et al., 2013) and the Mixed-Effected Model of Evolution (MEME). We also used the adaptive branch-site random effects likelihood (aBSREL) method (Smith et al., 2015) to test whether individual branches within the phylogeny experienced positive or a purifying selection. The FUBAR method uses a Bayesian approach to infer Ka/Ks substitution rates on a per-site basis, assuming that the strength of selection is constant across the phylogeny. This method may have power when positive selection is relatively weak (i.e. low values of ω>1). The MEME method, which employs a maximum likelihood approach (Murrell et al., 2012), tests sites subject to episodic positive selection under a proportion of branches, and positive selection for each site is inferred when positive beta is greater than alpha (β+ >α) and p-values are significant. Finally, aBSREL infers the optimal number of ω classes for each branch and then compares the full model to a null model in which ω < 1 across branches. All HyPhy methods were run under default parameters.

We used within PAML, the CodeML program to analyze the selective pressure of each codon based on six different maximum likelihood models via likelihood ratio tests (LRT) as previously described by Yang, 2007. Such models are: M0 (one ω ratio), M1a (nearly neutral), M2a (positive selection), M3 (discrete), M7 (beta) and M8 (beta & ω). We used the Bayes Empirical Bayes (BEB) method with a P > 95% threshold to determine evidence of positive selection for each site. Finally, we mapped into the phylogeny the amino acid changes for sites under positive selection, with the branch showing episodic positive selection and the elevation reported for each sample (see Table S1).

## RESULTS

### Mitogenomes: assembly and content

Our genome skimming approach generated 17 complete mitogenomes from nine species of NWJs. The number of initial input reads used for each assembly are presented in Table S2. The number of assembled reads in all approaches ranged from hundreds of thousands to three million. The low coverage (∼5X) samples showed the lowest number of assembled reads and the NOVOPlasty assembler showed the highest number of assembled reads used (Supplementary Information, Fig. S1a).

The organelle average coverage per assembly ranged from 18.7X to 31,148X (Supplementary Information, Fig. S1b). NOVOPlasty yielded higher average organelle coverage compared to the other assemblers (Supplementary Information, Fig. S1b), likely because it used the total available reads from the HTS data as input. Some mitogenomes circularized by AWA gave low average alignment scores below -3 (Supplementary Information, Table S3). After manually curating these sequences, we found unique insertions in some of the assemblies. The only notable differences found between the mitogenome assemblies from NOVOPlasty and MITObim/AWA were occasional insertions (size range 100-400 bp) found within some MITObim and AWA subset partitions assemblies of the same sample.

Circular mitogenomes assembled by NOVOPlasty showed less variation in length among species, ranging from 16,883 to 16,916 bp (Supplementary Information, Fig. S2a and Table S3). For the assemblies produced by MITObim, the length ranged from 17,158 to 19,385 bp and after removing overlapping sequences by AWA, the putative final length was between 16,886 and 18,578 bp (Supplementary Information, Fig. S2a and Table S3).

The percentage of mitochondrial DNA sequenced during HTS of whole-genomes, ranged between 0.1% and 2.29% for all assemblers. Samples with low sequencing coverage (∼5X) or preserved in ethanol had the lowest percentage of mtDNA, while samples flash-frozen or buffered directly in RNAlater had the highest percentage. Interestingly, the genomes with the lowest coverage showed a slightly higher percentage of mtDNA with NOVOPlasty compared with the other assemblers (Supplementary Information, Fig. S2b).

All nine species showed the standard avian gene order consisting of 13 PCGs, 22 tRNAs, 2 rRNAs, and one CR (Fig. 1). The majority of PCGs started with the Met (ATG) codon, except for COX1, which started with Val (GTG). We found the four typical types of stop codons (TAA, TAG, AGA, and AGG) in the mitogenomes. ND4 and COX3 had incomplete stop codons, as reported in other corvids (Liu et al., 2018) and parakeets (Dey et al., 2021). Additionally, gene couples, such as ATP8-ATP6 and ND4L-ND4, overlapped at their 3’ and 5’ ends, a phenomenon reported in metazoans (Shtolz & Mishmar 2023). With the exception of 8 tRNAs and ND6, which were located on the light strand, all other genes were located on the heavy strand.

**Fig. 1.**
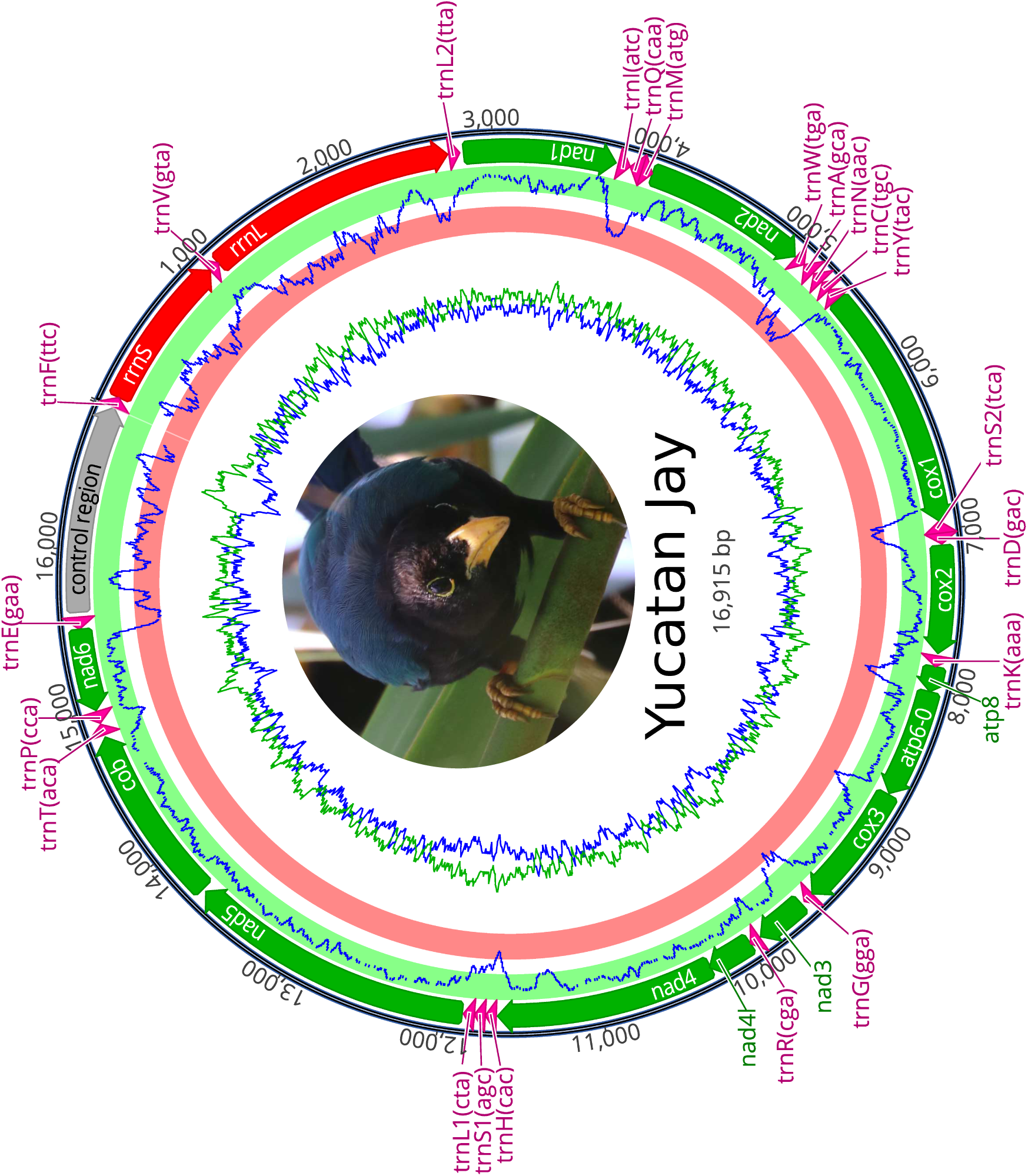
Mitogenome of the Yucatan jay (*Cyanocorax yucatanicus*) as a representative of New World Jay (NWJ) mitogenomes. The 16,916 bp mitogenome is depicted, with green annotations for protein-coding genes, red for rRNAs, pink for tRNAs, and gray for the control region. AT and GC content is represented by green and blue lines respectively in 20 bp windows. The green hoop indicates coding regions, while the red hoop represents non-coding regions.

The mitogenome base compositions varied slightly between species and even less at the intraespecific level (Fig. S3). AT and GC content percentages ranged from 55% to 56.5% and 42 to 44.5 %, respectively (Fig. S3). The AT and GC skews ranged from 0.085 to 0.11 and -0.32 to -0.35, respectively.

### Phylogenetic relationships of NWJ genera

All phylogenies based on the whole partitioned mitogenome, the CR and the PCGs agreed in topology (Supplementary Information, Fig. S4). In contrast, the phylogenies based on the rRNA genes only had unresolved relationships between the Azure-hooded jay, Steller’s jay, Unicolored jay and Pinyon jay (Supplementary Information, Fig. S4).

Overall pairwise genetic differences across the entire mitogenomes showed as expected the highest differences between the Eurasian magpie and all the NWJs ranging from 11.36% (Unicolored jay) to 12.24% (Black-throated magpie jay) (Supplementary Information, Fig. S5a). Within jays, the highest divergence was found between the Azure-hooded and Yucatan jays (11.5%), whereas the lowest was between the Yucatan and Bushy-crested jays (4.6%) (Supplementary Information, Fig. S5a). For downstream analyses, we used the phylogeny based on PCGs generated with MrBayes, using one sample per species (Supplementary Information, Fig. S5b).

Our results suggested a relationship between base composition and mitogenome size (Fig 2), while taking into account any possible dependence among species due to phylogenetic relatedness. Analysis with PGLS revealed a positive relationship between AT % and mitogenome length (Figure 2a & Table S4), whereas a negative relationship was found for GC % (Figure 2b & Table S4). The phylogenetic signal for AT % and mitogenome length was significant for both Brownian (t=2.84, pval=0.02) and OU models (t= 3.60, pval=0.00), whereas for GC % only the OU model (t=0.11, pval=0.04) showed marginal significance (see Table S4).

**Fig. 2.**
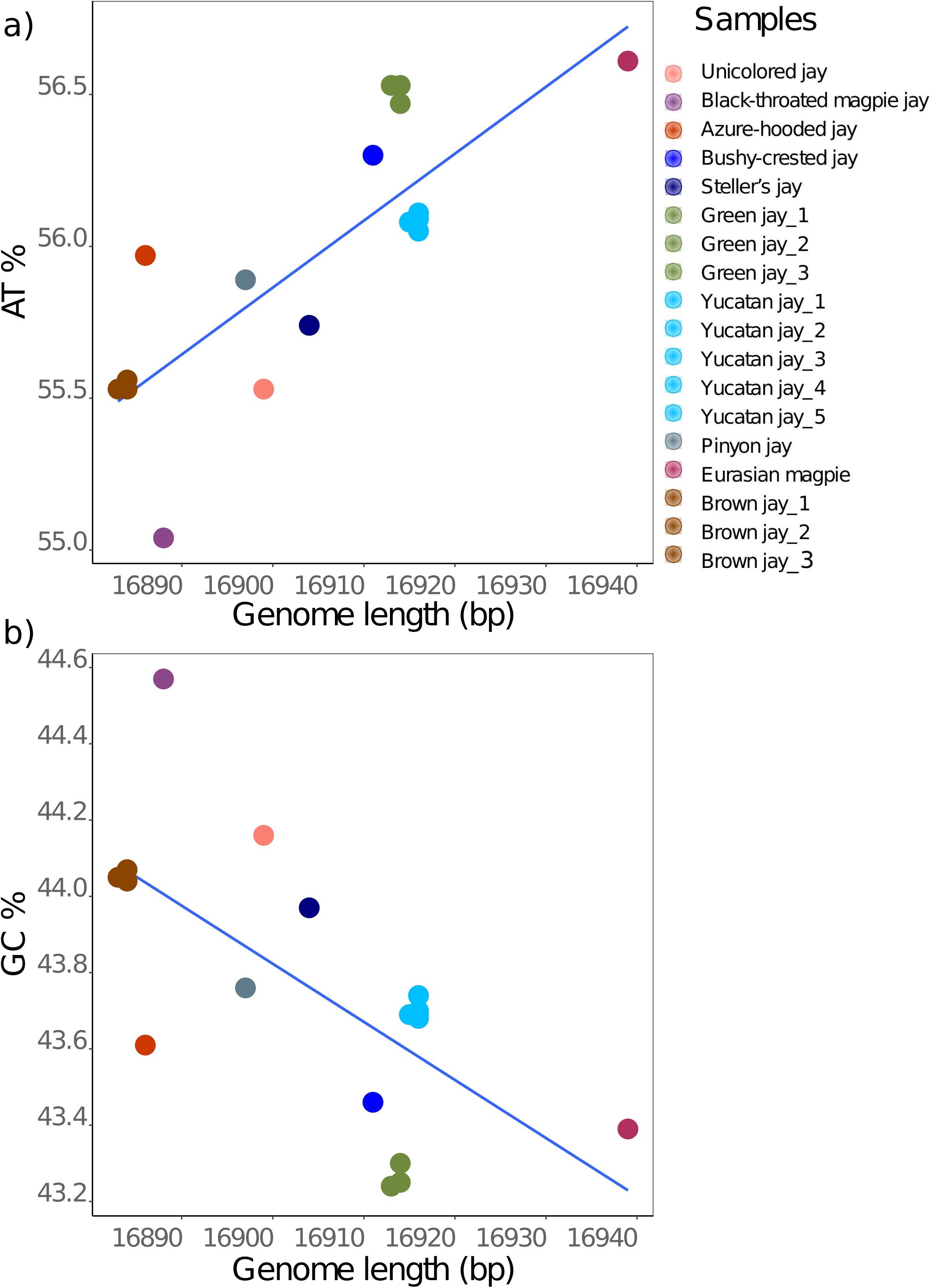
Scatterplots illustrating the relationship between genome length (bp) and base composition content (AT/GC) with a significant phylogenetic signal according to the Phylogenetic Generalized Least Squares (PGLS) analyses. a) PGLS regression model for genome length and AT content demonstrates a significant correlation (p = 0.0069, alpha = 2.5) under an OU model; b) PGLS regression model for genome length and GC content reveals a marginally significant correlation (p = 0.0449, alpha = 23.3) under an OU model. For additional information, refer to Table S4.

### Heteroplasmy: levels and locations

We explored levels of heteroplasmy in our assembled mitogenomes using NOVOPlasty, while increasing the amount of memory and, consequently, the number of reads used. Levels of heteroplasmy decreased with increasing number of used reads (Fig. S6). Most of the samples reached a stable number of heteoplasmic sites when 70% of the total reads were used (Fig. S6). However, for the three low-coverage genomes, NOVOPlasty always used the total reads available (Fig. S6). Regardless of the number of reads used, heteroplasmy levels were low in most samples, and the number of heteroplasmic sites ranged from 0 to 7, except for Unicoloured Jay which had 21 sites (Fig. S6).

The alternative allele frequencies of the identified heteroplasmic sites ranged from 0.005 to 0.5 (Fig. 3). The highest allele frequencies were found within the CR of all mitogenomes followed by sites located in the mitoribosome rrnL of the Azure-hooded jay and the Unicolored jay (Fig. 3). For PCGs, heteroplasmic sites in COX1, NAD2, and NAD5 were found for three species, with a minor allele frequency up to 0.13 (see Fig. 3). The Unicolored jay had the lowest heteroplasmic allele frequencies while containing the highest number of heteroplasmic sites among the NWJs (Figs. 3 and 4).

**Fig. 3.**
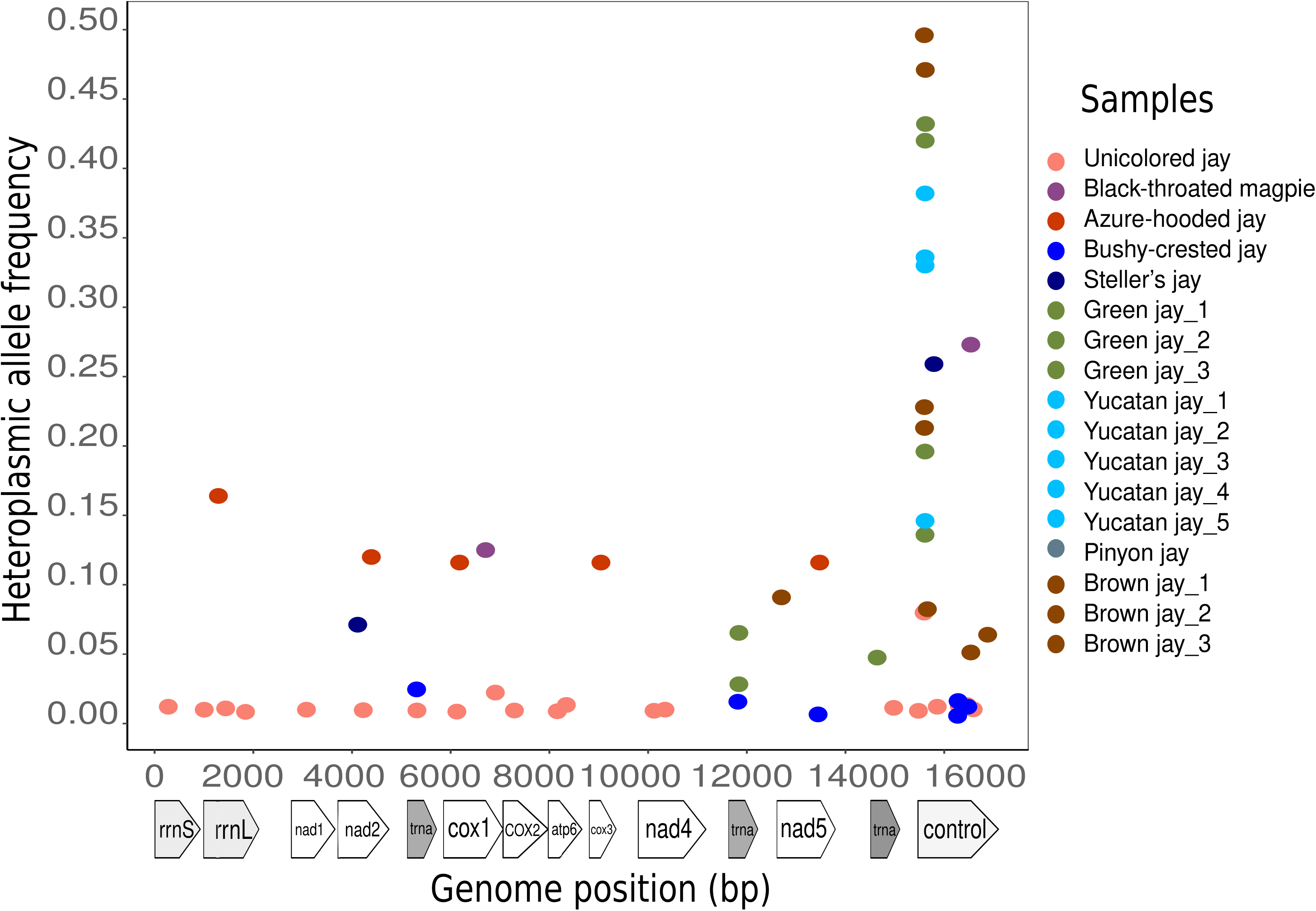
Scatterplot depicting heteroplasmic allele frequencies and their genome position across all NWJ mitogenomes. The Unicolored jay exhibits the highest heteroplasmy levels (21 sites) but with comparatively lower allele frequencies, while the beginning of the control region corresponds to the highest value in several NWJs.

**Fig. 4.**
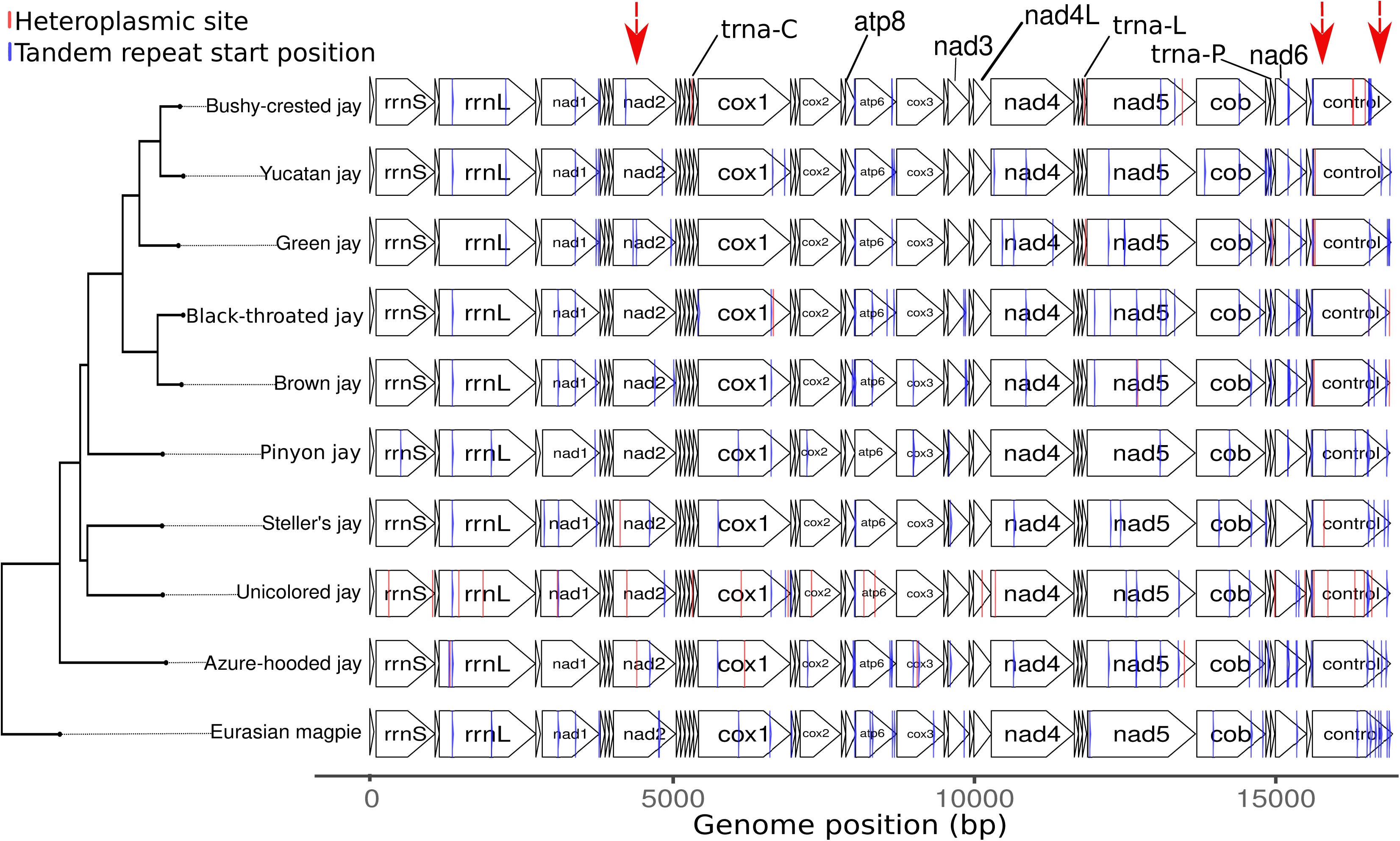
Locations of tandem repeats (TRs, blue lines) and heteroplasmic sites (red lines) in NWJ mitogenomes, with the Eurasian magpie as the outgroup. The figure highlights specific patterns using red arrows. In the control region, the number of tandem repeats decreases across the phylogeny. Another arrow indicates co-occurrence of heteroplasmic sites and tandem repeats at the same site. Additionally, the arrow in gene ND2 demonstrates heteroplasmic sites and tandem repeats showing divergence throughout the phylogeny. The Eurasian magpie has no data available for heteroplasmy analysis.

Most heteroplasmic sites within the CR were in low complexity regions or TRs (see alignments in Supplementary Information, Fig. S7a). Single nucleotide repeats of either Cs or Ts were located at the beginning of the CR (domain I), an area of the mitogenome with high variability. The position of heteroplasmic sites varied between species, but in closely related species some sites were in the same position or in close proximity (Fig. 4). For example, one heteroplasmic site at the beginning of the CR was located in the same position (or one base pair away) in the *Cyanocorax* lineage (Yucatan, Green and Bushy-crested jays) and in the Brown and Unicolored jays (see alignments in Fig. S7a). We found heteroplasmic sites in genes NAD2, NAD5 and COX1 in close proximity (∼57 - 300 bases apart) between the Azure-hooded jay and the Steller’s and Unicolored jays (Fig. 4; see alignments in Fig. S7b and Fig. S8a-b). Another interesting heteroplasmic site was present on tRNA-L and located 20 bases apart between the Green and Bushy-crested jays (Fig. 4; see alignments in Fig. S8c).

A single putative NUMT, measuring 125 base pairs in length, was detected in the Bushy-crested jay. It was aligned at the end of the CR. Notably, the NUMT emerged when we used only 10 to 30% of the total reads.

### Tandem repeats: features and location

We obtained a general overview of the repeat composition in the mitogenomes using Tandem Repeats Finder after the multiple alignment. Additionally, we used the phylogenetic context to identify some evolutionary trends in TRs. First, we found medium entropy values (metric for nucleotide composition) for each sample ranging from 1.3 up to 1.6 per sample (Fig. S9a), where a value of 2 indicates a conserved TR (little variation in nucleotide). The Green jay shows the lowest entropy values, and the Steller’s and Unicolored jays the highest value (Fig. S9a). Second, we identify that the total number and mean length of TRs varied per genomic region (Fig. 5) and species. Overall, the TRs in the NWJ mitogenomes comprised more than 450 bp, this is up to 3% of their mitogenomes (Fig. S9b), where species with fewer TRs tend to have longer TRs (Fig. S9c).

**Fig. 5.**
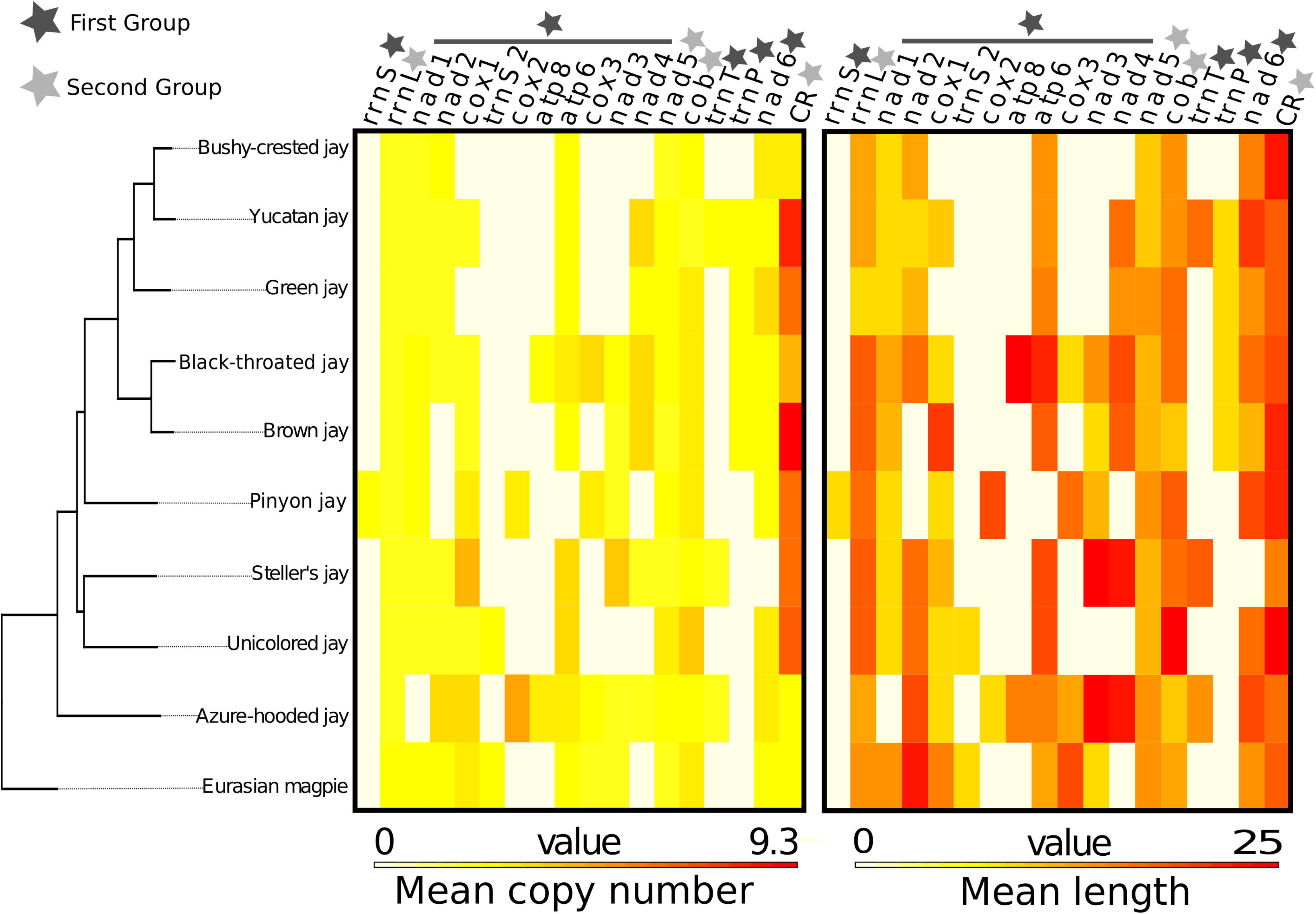
Heatmaps comparing copy number (tandem unit) and length (bp) of tandem repeats (TRs) in NWJ mitogenomes, with the Eurasian magpie as the outgroup. The heatmaps display mitochondrial regions ordered by genome position. Black stars indicate regions from the first group where TRs are not present in all species, while gray stars indicate regions from the second group where TRs are present in all species.

The ancestral reconstruction of the structure of TRs showed trends for both increases and decreases in length, but also losses and gains in the overall number of TRs across the phylogeny. We identified two sets of mitochondrial regions exhibiting different TR trajectories. The first set is characterized by TRs that were unique to individual species and comprised most of the PCGs, mitoribosomes and transfer RNAs (see black stars in Fig. 5). The second set is characterized by TRs contained in all species, this includes the CR, two genes and one mitoribosome (see gray stars in Fig. 5)

In the first set most genes followed dynamic trajectories in TR lengths and copy number. However, the TR trajectories of transfer RNAs were unique. For example, the unique TR in the trnT (up to 18 nucleotides long) is gained in just three lineages (Figs. 4 and 5). Likewise, in the trnP a unique TR (short tandem repeat of 9 nucleotides long) is gained in a clade with the *Cyanocorax* lineage (Yucatan and Green jays), the Brown and the Black-throated magpie jays (except for the Bushy-crested jay) (Figs. 4 and 5). We hypothesized that the TR gained in the trnP was associated with low elevation after examining the ancestral reconstruction (Fig. 6a and b). Analysis with PGLS revealed a significant association (r=-0.443, t=-7.143126, pval=0.0001) between elevation and the presence of the TR in trnP, with high phylogenetic signal (see Fig. 6b and Table S4).

**Fig. 6.**
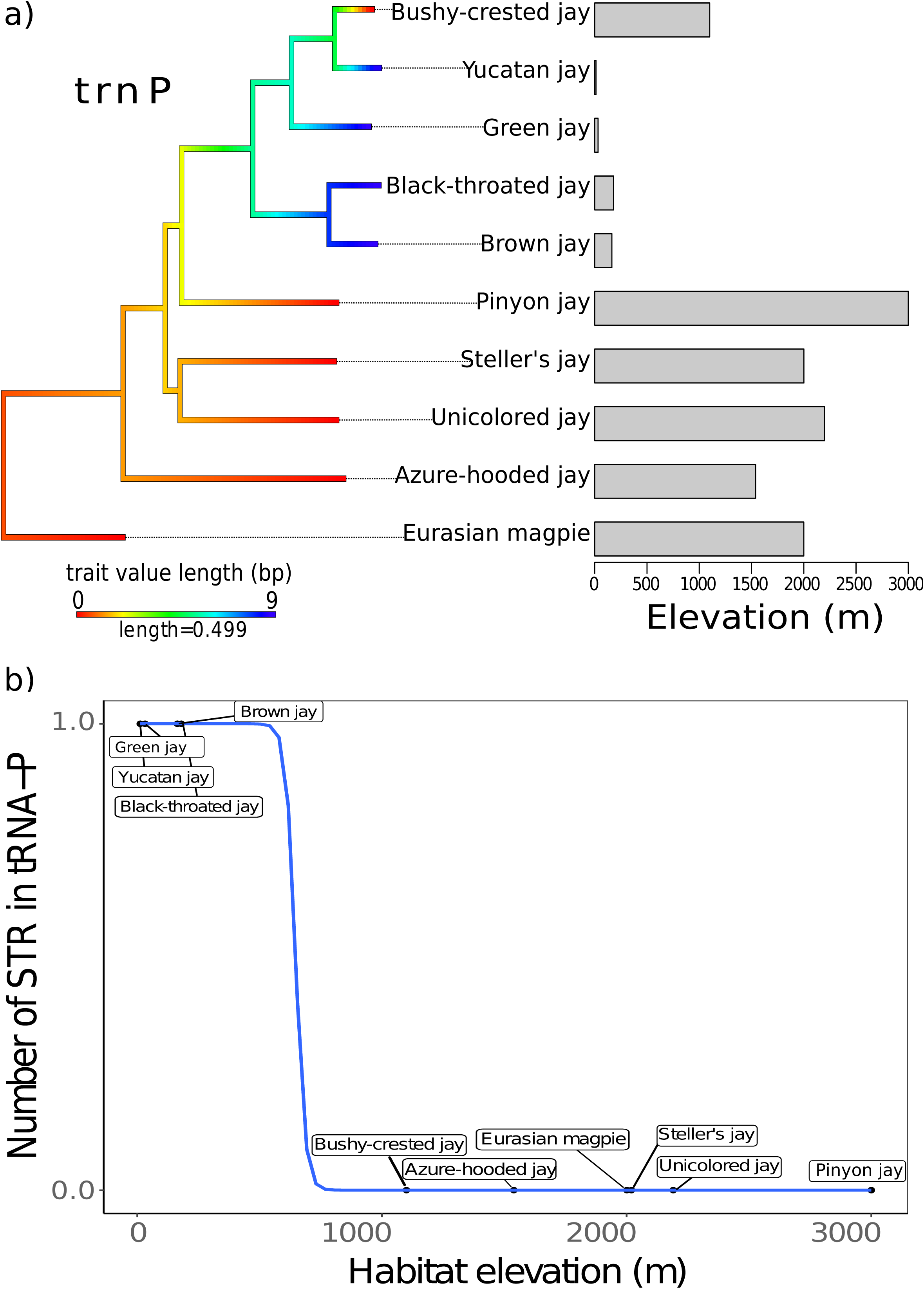
Comparison of habitat elevation with the length and total number of a TR in tRNA-P. a) Ancestral state reconstruction of the TR length in tRNA-P for NWJs and the Eurasian magpie as the outgroup. The contour map phylogeny displays the estimated evolutionary history, while the bar plot indicates that lowland jays have a 9 bp long TR; b) Binomial regression demonstrates the association between TR presence/absence in tRNA-P and habitat elevation in NWJs, where the PGLS analysis show significant correlation (p = 0.0001, alpha = 5.09, see details in Table S4).

The second mitochondrial set, showed a loss of TRs on the phylogeny (except for NAD5) accompanied by an elongation. For example, over some paths of the evolutionary history of the NWJs, the number of TRs in the CR went from 8 in the NWJ common ancestor to 3 TRs in the Busy-crested Jay (Fig. 4), while the mean length increased slightly in most lineages (between 10 to 24 nucleotides; Fig. 5). The mitoribosome rrnL showed an elongated TR in basal species becoming smaller in the *Cyanocorax* lineage (Fig. 5).

### Selection in mitogenomes

The analyses performed for natural selection such as the site-by-site and branch model approaches suggest an overall patter of purifying selection, with five sites showing statistically significant evidence for positive selection. Estimates with MEGA showed ω < 1 for each PCG, suggesting an overall pattern of purifying selection (Supplementary Information, Fig. S10a). ATP8 had the highest ω value (0.13), whereas COX1 had the lowest (0.01) (Fig. S10a). More than 60% of the codons in all PCGs are under purifying selection (Fig. S10b). Supporting these results, the FUBAR method also found evidence of pervasive negative selection on 30% of total codons (3,347 codon sites) across all mitochondrial PCGs.

Model comparisons implemented in PAML rejected the hypothesis that evolution of the mitogenomes is neutral, and suggested that ω varies among sites (pval < 0.001) (see Table S6). Within the preferred alternative model (M8), two codons showed posterior probability scores > 0.9 according to the Bayes Empirical Bayes (BEB) approach (Yang et al., 2007) and were located within COX2 and ND2 (Table S5 & S6).

A total of 16 sites across all 13 PCGs showed significant levels of positive selection based on the HyPhy (FUBAR & MEME) or PAML (CodeML) (Fig. 7 & Table S6). We detected five sites with statistically significant evidence for positive selection with two or three methods (see Table S6 & Fig. 8). We mapped the amino acids changes at these five sites with the phylogeny and found that the changes were mainly present in the species living at low elevations (Fig. 8).

**Fig. 7.**
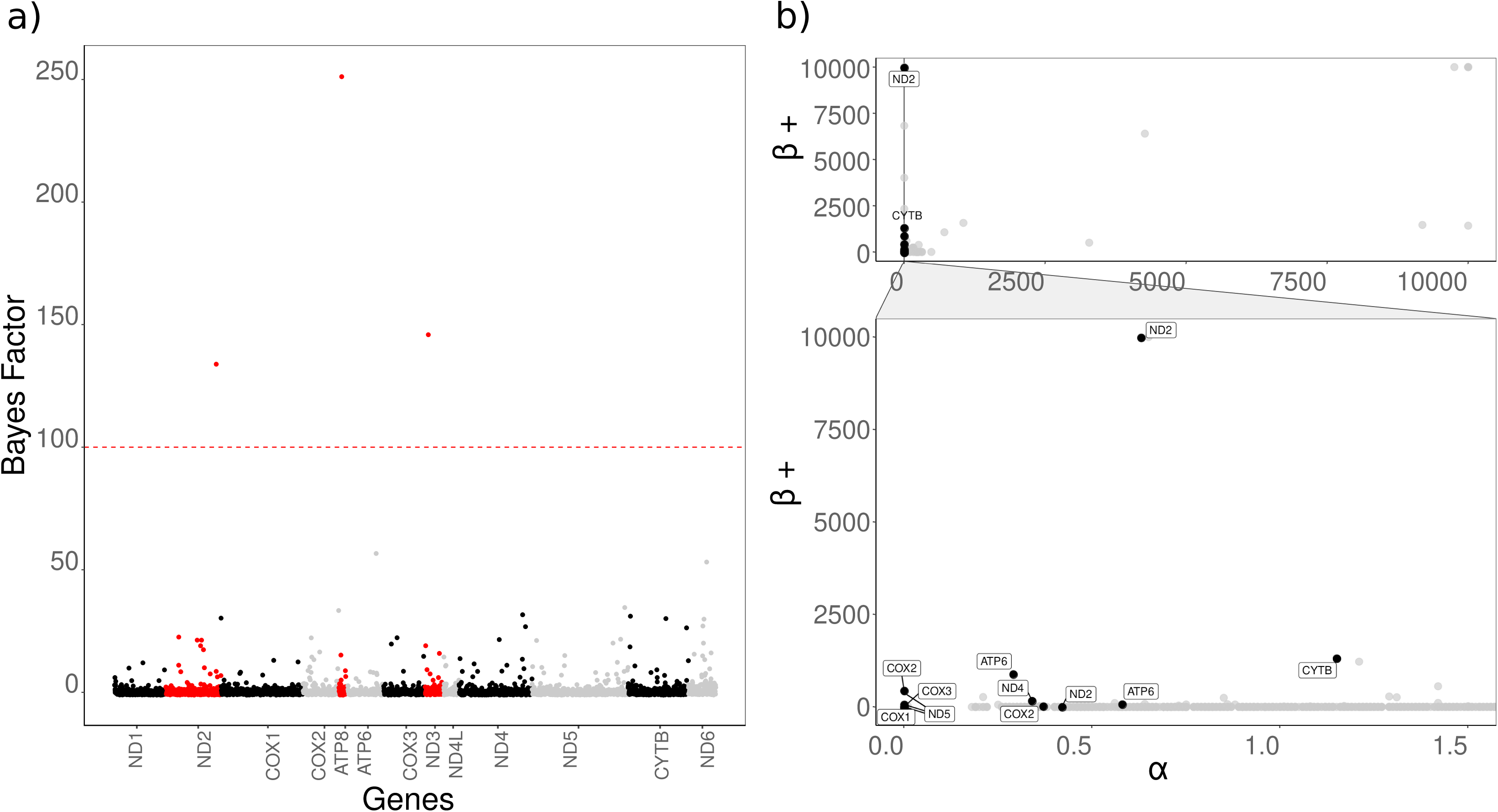
Selection analyses in protein-coding genes (PCG) across NWJs mitogenomes. a) Manhattan-like plot displays the Bayes factor for each site, with red showing genes and sites where Bayes factor > 100 and posterior probabilities > 0.9, indicating strong evidence for positive selection using the Fast Unconstrained Bayesian Approximation (FUBAR) analysis. The red dashed line represents the threshold for strong and decisive evidence of positive selection; b) Scatterplot with a zoomed-in view (gray triangle) showcases positive beta and alpha values. Black dots represent sites under positive selection with a p-value threshold of 0.05, based on the Mixed-Effected Model of Evolution (MEME) analysis.

**Fig. 8.**
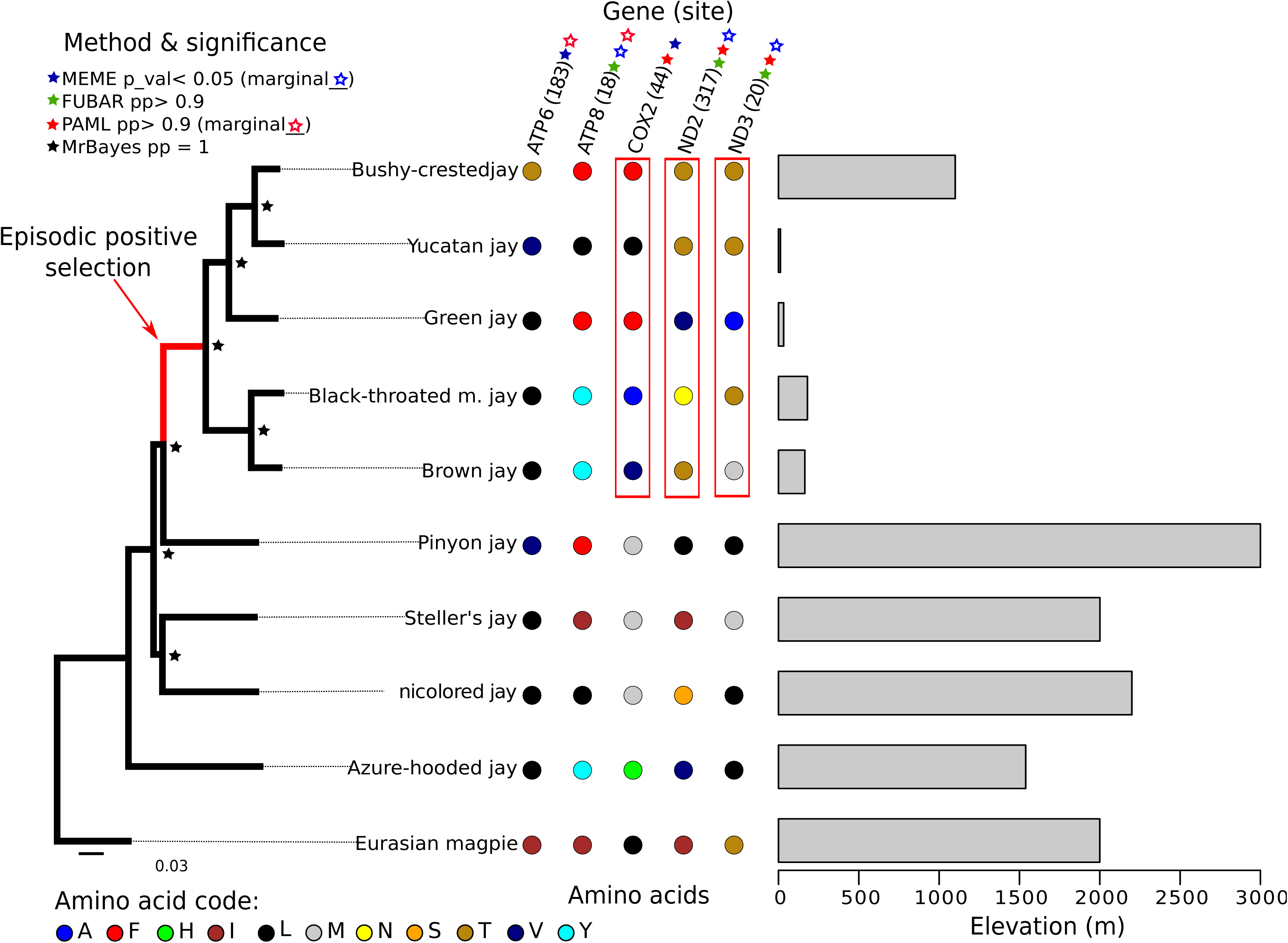
Linking habitat elevation with sites under positive selection and amino acid changes in NWJ mitogenomes. The branch undergoing episodic positive selection, identified by the adaptive branch-site random effects likelihood (aBSREL) method, is highlighted in red. Phylogenetic support is represented by posterior probabilities from MrBayes. Amino acid changes in the five selected sites, supported by at least two of the three methods used, are indicated by dotted colors. A filled-out star denotes a statistically significant site, while an outlined star indicates marginal statistical support. The bars depict the relative habitat elevation in meters for each species.

Finally, the aBSREL method found evidence of episodic diversifying selection (corrected p-value = 0.037) occurring on one branch of the NWJ phylogeny (Fig. 8). This branch leads to clades undergoing a transition in habitat from highlands to lowlands (see Fig. 8).

## DISCUSSION

We present our findings on the mitogenomes of nine species of NWJ’s, obtained by skimming genomes from HTS using multiple bioinformatic tools. We analyzed the NWJ mitogenomes and found evidence of molecular changes that occurred during adaptation to low elevation. Our study yielded four key observations: Firstly, heteroplasmic sites in NWJ mitogenomes are concentrated in low complex regions or TRs, suggesting that TRs may be a target for heteroplasmic mutations, and possibly regulation and/or recombination may be linked to these regions. Secondly, we identified a TR in a transfer RNA that was present in a clade associated with changes in habitat elevation. Thirdly, we found that TRs are dynamic and vary across mitochondrial regions, with TRs in the CR elongating while reducing in amount in the NWJs evolutionary history. Finally, our analysis revealed changes in amino acids in some OXPHOS genes also associated with habitat elevation. Our findings provide insights into the adaptive mechanisms of NWJ to low elevation and shed light on the dynamic nature of their mitogenomes.

### NWJ mitogenomes

As expected for the Corvidae family, the assembled NWJ mitogenomes contain the typical 37 mitochondrial genes in standard avian order (Krzeminska et al., 2016; Li et al., 2016; Johnsen et al., 2017; Sarker et al., 2017) with similar features as in other metazoans (Shtolz & Mishmar 2023). Our results confirm that gene arrangements are stable within taxonomic groups with a few exceptions, whereas variations occur between major groups (Boore, 1999).

Assembling the mitogenomes with different approaches allowed us to compare and assess the consistency of the mitogenome assemblies. MITObim and AWA produced high-quality scores with some inconsistent regions. The mitogenomes from NOVOPlasty did not show any inserted regions within the PCGs and used all the available reads for the assembly. Hence, the mitogenomes generated with NOVOPlasty were chosen as the final assemblies. A shortcoming of our study is that we were unable to detect long TRs, duplicated genes or CRs because we used HTS. We know that LRS has been effective in detecting complex mitogenome organization (Macey et al., 2021; Vertebrate Genomes Project Consortium et al., 2021). Nonetheless, the gene arrangement in the NWJ mitogenomes is consistent with that of other Corvidae species (Krzeminska et al., 2016). Therefore, our final mitogenomes are robust for comparative studies.

The amount of mitochondrial DNA recovered from HTS of whole-genomes was less than 3% of the total data, which is influenced by several factors such as sequencing coverage, tissue type and preservation conditions. Our samples preserved in ethanol recovered lower mtDNA compared to flash-frozen and buffered in RNAlater samples. The location of the muscle tissue may also play a role. For example, we only know that vertebrate muscle tissue tends to yield more mtDNA due to its high metabolic activity and phosphorylation levels (Butler, 2016; Kovacs & Meyers, 2000). However, pectoral versus leg muscle may have differences in metabolic rate and in the amount of mtDNA, thus, further research is needed to explore this topic.

The mitogenome base composition is reported to be influenced by gene function and regulation (Li & Du, 2014). Base composition between the NWJs varies slightly among species and is a metric that distinguishes them. We found that the NWJ mitogenomes base composition was skewed towards AT, consistent with observations in the Yellow-billed babblers (Wang et al., 2015) and the weasels (Skorupski, 2022). Our finding supports the hypothesis that codon usage bias is positively correlated with the AT content in the third codon position (Skorupski, 2022). In addition, we found a trend between the mitogenome size and base composition. The trend suggests an increase in mitogenome length accompanied by an increase in AT but a decrease in GC. This relationship had a phylogenetic signal following an OU model of evolution and could be associated not only with the codon usage but also with AT rich regions, such as in arthropods (Fauron, & Wolstenholme 1976; Zhou et al., 2007), and the TRs dynamics. It is known that the sequence expansions in animal mitogenomes are associated to either non-coding unique sequences or repeats in the CR (Wolstenholme, 1992). For example, the AT rich regions in *Drosophila* species located mostly in the CR, contains the origin of replication and changes more rapidly than the rest of the mitogenome (Shah & Langley 1979). Further investigation is needed to fully understand AT rich regions in birds, its implications in replications and mitogenome size.

### Mitochondrial phylogenetic relationships in the NWJs

We inferred phylogenetic trees based on the whole partitioned mitogenome with MrBayes. The PCGs and the CR agreed on topology with high levels of support for both datasets. The results overall agreed with previous studies based on other genetic markers (Ericson et al., 2005; Bonaccorso & Townsend Peterson, 2007; Bonaccorso et al., 2010; Fernando et al., 2017; Saunders & Edwards, 2000). Our results confirm that the CR is a useful phylogenetic marker for NWJs and that the tree topology agrees with previous results based on Saunders and Edwards (2000), but differs from others solely regarding the placement of the Pinyon jay (Bonaccorso & Townsend Peterson, 2007; Fernando et al., 2017). It is likely that the discordance in the placement of the Pinyon jay is due to the differences in the composition of TRs that we found in this species. In addition, the mitoribosomes dataset was less able to resolve phylogenetic inference compared to other datasets, likely driven by the content of species-specific TRs (shown in this study) and mitonuclear coevolution as in mammals (see explanation below) (Weaver et al., 2022).

### Detecting heteroplasmy from HTS data and NUMTs

The genetic variation of mitochondrial DNA at the individual level, known as heteroplasmy, has shed valuable insights on mitogenome evolutionary and cellular processes. Heteroplasmy can result from either mutational mechanisms or a paternal leak of mtDNA (Ingman & Gyllensten, 2006; Ye et al., 2022). With older techniques, the accurate detection of heteroplasmy was difficult to achieve because NUMTs mimic polymorphism (Parakatselaki et al., 2022). However, considerable levels of heteroplasmy have been reported in nematodes (*C. elegans,* by Konrad et al., 2017), crabs (Koolkarnkhai et al., 2019; Rodríguez-Pena et al., 2020), birds such as the partridge (Gandolfi et al., 2017; Pizzirani et al., 2020) and the Crested ibis (He et al., 2013), mammals (Burgstaller et al., 2018), turtles (Tikochinski et al., 2020) and the Tuatara (Macey et al., 2021). With the increased availability of HTS and LRS in growing numbers of taxa, heteroplasmy is becoming more easily detectable and is more likely to be the rule rather than the exception (reviewed in Pereira et al., 2021).

Detecting heteroplasmy levels from HTS requires the consideration of many factors, such as tissue type, preservation conditions, sequencing coverage and type of analyses. There is tissue-specific evolution of polymorphism in heteroplasmic animals, which can bias the results (Casane et al., 1994; Casane & Guéride, 2002). Thus, to obtain confident heteroplasmy levels, one should ideally use the same tissue type between samples, the same preservation conditions and sequencing coverage. By performing multiple heteroplasmy analyses on muscle tissue, we were able to achieve confident heteroplasmy levels. During the analyses we found it relevant to address the saturation point of heteroplasmy levels, according to the data type of each sample. The final levels of heteroplasmy detected in the NWJ mitogenomes varied according to the sequencing coverage and the initial amount of reads used. For this reason, we recommend an initial 25X sequencing coverage for genome skimming and heteroplasmy detection. We also recommend using up to 70% of the total reads to get the appropriate organelle coverage and to use an MAF threshold of 0.01.

NUMTs have a major impact on mitogenomes and they may confound cases of heteroplasmy (Parakatselaki & Ladoukakis, 2021; Parakatselaki et al., 2022). We used the latest version of NOVOPlasty that performed additional re-alignments, phasing and thresholds (in length and mutations) to distinguish between NUMTs and heteroplasmy. We detected only one possible NUMT in the Bushy-crested jay with a low subset of reads (10 – 30%). Due to the low confidence of this unique NUMT, we note that for the detection, curation and analysis of real NUMTs, more examination is required. Thus, NUMTs in the NWJs must remain a topic for future investigation. A step to follow would be to align the mtDNA with the whole nuclear genome, making NUMTs another area of mitogenomics where we expect to see rapid development (Zardoya, 2020).

### High levels of heteroplasmy in the Unicolored jay

Heteroplasmy levels in the NWJs were relatively low compared to the chicken (up to 178 sites, Huang et al., 2019) and turtles (up to 300 sites, Tikochinski et al., 2020). In the NWJs, some samples were homoplasmic while the Unicolored jay showed higher heteroplasmy levels than the other NWJs. This particular sample may have been a highly heteroplasmic individual. Recent studies have demonstrated that heteroplasmy composition at the individual level is a population signature and, at the population level, provides a unique fingerprint of family clustering (Tikochinski et al., 2020).

The biology of the Unicolored jay may contribute to its higher levels of heteroplasmy compared to the other NWJs. This species may have experienced population bottlenecks due to its limited dispersion, and isolation in small cloud forest patches (Webber & Brown, 1994). According to Venkatraman and collaborators (2019), the patches became isolated after the Pleistocene, with glacial cycles driving speciation up to five lineages (at the genetic and niche level). Also, climate change and land use have had a significant impact on the cloud forest (Lawton et al., 2001; Helmer et al., 2019), exacerbating the isolation of the Unicolored jay population. For example, the lineage Unicolored jay (*guerrerensis*) -- our Unicolored jay sample – is limited to a very small patch of cloud forest (∼ 100 km x 40km) (McCormack et al., 2008). Finally, hybridization with other Mexican Jays cannot be ruled out, as the Unicolored jay lineage has not been found to be reproductively isolated, and is sympatric to other Mexican Jays (McCormack et al., 2008; 2009). High heteroplasmic levels are reported in hybrids, such as between the Rock partridge (*Alectoris graeca*) and the Chukar partridge (*Alectoris chukar*), with potential paternal leakage (Gandolfi et al., 2017). Unfortunately, we have neither pedigrees to test for paternal linkage nor samples from sympatric species to test for hybridization in the Unicolored jay.

Tikochinski and collaborators (2020) reported a trend in turtles similar to the one we found in the Unicolored jay, such as higher heteroplasmy levels likely driven by a combination of high phylopatry and habitat reduction. They also demonstrated that, in natural populations, heteroplasmy might accumulate and help maintain genetic diversity during a sudden decline in population size and add population structure during the recovery. For instance, our results may also suggest that heteroplasmy levels in the Unicolored jay must be driven by repeated bottlenecks, in combination to the cellular bottleneck during gametogenesis. Furthermore, purifying selection with drift might be one of the major ways that heteroplasmic mutations with deleterious effects are eradicated (Konrad et al., 2017; Koolkarnkhai et al., 2019). This explains the low allele frequencies of heteroplasmy mutation in the Unicolored jay compared to the other NWJs, likely driven by drift and purifying selection.

Despite how little we know about heteroplasmy, its detection with HTS has proven to be effective (Huang et al., 2019), even for detecting low-frequency mutations (Li et al., 2010; Konrad et al., 2017). We notice a promising research avenue in the discussion of the function and evolution of heteroplasmy in natural populations. It is likely a potential molecular marker in the fields of wildlife management, conservation genetics and biodiversity studies (Tikochinski et al., 2020).

### Location of heteroplasmic sites in TRs, in the CR and genome-wide

Numerous studies have reported the presence of heteroplasmic sites in the CR and tandem repeats (TRs) of mitochondrial genomes across various taxa (reviewed in Lunt et al., 1998; Casane & Guéride, 2002; He et al., 2013). For example, in invertebrates (Koolkarnkhai et al., 2019; Ye et al., 2022) and vertebrates (Vertebrate Genomes Project Consortium et al., 2021), such as in humans (Li et al., 2010), birds (Mundy et al., 1996b; Huang et al., 2019; Torres et al., 2019; Pizzirani et al., 2020) and reptiles (Ray, 2003; Tikochinski et al., 2020). We found that the CR has the highest number of heteroplasmic sites in most NWJ species, particularly in the domain I region known for its variability (Abbott et al., 2005; Saunders & Edwards, 2000). Indeed, studies suggest that heteroplasmic mutations in TRs within the CR are useful for studying repeat duplication (Mundy et al., 1996b; Mundy & Helbig, 2004).

Heteroplasmic mutations seems to act differently throughout the mitogenome of the NWJs. The allele frequencies of heteroplasmic sites in the CR of the NWJs are generally higher than the rest of the mitogenomes. Cases with a high allele frequency of heteroplasmic mutations could be promoted by selection, where containing more than one mitochondrial haplotype may offer an advantage (Ingman & Gyllensten, 2006; Li et al., 2010; Tikochinski et al., 2020; Macey et al., 2021). We know so far that in cows, de novo heteroplasmic mutations have been fixed in two or three generations without compromising fitness (Olivo et al., 1983). On the other hand, the heteroplasmic sites in PCGs and RNAs were often located in or near TRs with lower allele frequencies than the CR, suggesting that purifying selection is stronger on PCGs. Furthermore, TRs in the CR are likely more susceptible to heteroplasmic mutations with a potential regulatory mechanism.

The landscape and degree of recombination in mitogenomes it still a matter of debate (Thapana et al., 2022; Ye et al., 2022), with some studies on vertebrates suggesting a role in the evolution of duplicated regions. In hornbill birds, a long TR duplication with heteroplasmy showed that the two duplicated regions evolved in concert and likely originated from recombination (Sammler et al., 2011). In varanids (*Varanus salvator*), the evolutionary pattern of duplicated CRs suggested that recombination events occured between the two CRs (Thapana et al., 2022). Finally, in the mitogenomes of seven bushtits (*Aegithalos*), recombination (at every replication cycle) was detected between specific regions of the duplicated CRs that evolved also in concert (Wang et al., 2015).

We did not find evidence for duplicated genes or CRs, but we have suggestive results for recombination in the NWJ mitogenomes. For example, the heteroplasmic site in a TR is in the same position in sister species but in basal linages is shifted from the TRs ∼300bp apart (in rrnL and NAD2). These patterns indicate other mechanisms might shift the heteroplasmic site besides the classical “slipped-strand mispairing”. Recombination likely occurs in an organized manner, potentially in TRs and in a specific section of the CR (Wang et al., 2015; Wang, Liu et al., 2015), particularly the beginning of domain I. Although, if recombination acts with paternal leakage -- alleged to be rare – the results will be hard to detect. Further research is needed to understand the mechanisms involved in heteroplasmic mutations and TRs in maintaining biogenesis, evolution, and function in the CR, which regulates transcription and replication (Kinkar et al., 2021).

### TRs in all species

TRs in vertebrate mitogenomes, such as in birds (*Ardea* sp., Zhou et al., 2014b) and certain fishes (*Symphurus* sp., Shi et al., 2015), are reported to be involved in long duplication and gene rearrangement following concerted evolution. The presence of TRs in the NWJ mitogenomes is highly variable. Some TRs have become shorter over time, while others have increased in length or have originated multiple times or within specific lineages. The TRs found in NAD and COX genes, were the most dynamic compared to other PCGs, potentially due to their role in the OXPHOS pathway. In contrast, the CR contains the majority of TRs, but their number decreased while elongated throughout the evolutionary history of NWJs.

TRs have a notable tendency to undergo elongation, irrespective of their origin. Over the course of evolutionary history, they often undergo multiple cycles of reduction or elimination in NWJs. The increase in copy numbers and length of TRs may be influenced by a combination of mutation, selection, and drift, as proposed in other vertebrates (Ba et al., 2016). However, most TRs undergo a constant process called “slipped-strand mispairing,” which promotes repetitive DNA and expands the TRs (Levinson & Gutman, 1987). Previous studies in the Yellow-Browed tit (*Sylviparus modestus*) demonstrated how TRs in the CR and other mitochondrial regions have multiple origins due to constant “slipped-strand mispairing” but when a long TR is in play (117bl long) this can originate from recombination (Wang et al., 2015). Several studies have shown that recombination may play an important role in long TR duplication (Sammler et al., 2011; Wang et al., 2015; Thapana et al., 2022; Ye et al., 2022).

Mitochondrial TRs likely impact phenotypes due to mitochondrial efficiency (Gemayel et al., 2012), similar to TRs in nuclear genomes that are associated to morphological plasticity (Fondon & Garner, 2004) and accelerate the evolution of gene expression (Usdin, 2008; Gemayel et al., 2010, 2012; Adrion et al., 2016).

Recent studies with LRS of mitogenomes have revealed that long TRs in non-coding regions regulate replication and/or transcription (Kinkar et al., 2021). In fact, a long TR (4.5 kb) in a population of the trematode (*Clonorchis sinensis*) plays a functional role in regulating replication and transcription, similar to the CR in mammals (Kinkar et al., 2020). This supports the hypothesis that long TRs may contribute to the origin of the CR (Kinkar et al., 2020), with evidence in birds having the ancestral state of a duplicated CR (Urantówka et al., 2020). Further research is needed to understand the impact of TRs in the mitogenome, their roles in coding and non-coding regions, and their contribution to the dynamics of morphological plasticity and other biological processes.

### Species-specific TRs and habitat elevation

We reported species-specific TRs in tRNAs in the NWJ mitogenomes. Interestingly, tRNAs in mitogenomes interact with the nuclear genome for mitochondrial protein translation (review by Karakaidos & Rampias, 2020). This interaction throughout gene expression in the nuclear genome leads to alterations in amino acid substitution rates in proteins that interact with mitogenome tRNAs (Adrion et al., 2016). Consequently, mitogenome tRNAs have higher mutation rates compared to other parts of mitogenomes and even nuclear tRNAs, but also undergo significant modifications during animal evolution (Helm et al., 2000; Tsutomu et al., 2011; Adrion et al., 2016). The mutational flexibility in tRNAs is likely an adaptation to increased mutation pressure associated with the evolution of animal complexity (Kuhle et al., 2020). Thus, the presence of species-specific TRs in mitochondrial tRNAs is likely a key to understand the interaction with the nuclear genome.

The most remarkable TR in a transfer RNA in the NWJs mitogenomes is present in tRNA-P (9 bp long), and it is specific to a clade and significantly associated with habitat elevation. This TR might confer some advantage to species found at low elevations. Montelli et al. (2015) reported similar findings in cetaceans, demonstrating that the evolution of mitochondrial tRNAs, indicated by changes in base composition of AT and GC-skew, is influenced by environmental factors in toothed whales living in freshwater and which are deep divers. We suspect that the other transfer RNAs – those with species-specific TRs, gained or lost multiple times in the NWJs -- are associated with other NWJ features beyond habitat elevation not captured in our study.

Further investigation is needed to expose the underlying mechanism driving TRs in the tRNA-P to facilitate the interaction with the modified proteins from the nuclear genome. The expression of the specific nuclear gene associated with adaptation to low elevation in the NWJs could provide some information about how and what protein modification interacts with the mitogenome tRNA-P.

### Positive selection in OXPHOS genes and habitat elevation

Purifying selection is the primary force shaping NWJ mitogenomes, particularly on the OXPHOS pathway essential for ATP and heat production (Meiklejohn et al., 2007; Morales et al., 2015). Here, selective pressure is high, emphasizing the functional integrity of proteins involved (Meiklejohn et al., 2007; Morales et al., 2015). Strong purifying selection in the mitogenome complex IV has been found in other avian studies, especially for the COX1 gene (Kerr, 2011; Ramos et al., 2018). In the NWJs, the COX1 gene presented the lowest ω value, pointing to high purifying selective pressures. This is expected since complex IV modulates protein configuration and adaptation, enhancing the coupling efficiency for ATP or heat production (Kerr, 2011; Ramos et al., 2018).

On the other hand, we also found signals for positive selection in five sites in the NWJ mitogenomes within complexes I, IV and V. Studies have found that, non-synonymous substitutions in complex I affect proton transport and the balance between energy and heat production (Garvin et al., 2015); while complex V is the final step of the OXPHOS pathway and synthesizes ATP using a proton gradient (Ramos et al., 2018). Thus, environmental conditions, like temperature and altitude, impose selective pressures on ATP and heat production trade-offs. Signatures of positive selection within genes of complex I have been found in avian species (Zink, 2005; Zhou et al., 2014; Lamb et al., 2018; Ramos et al., 2018) and other taxa (Garvin et al., 2015; Wang et al., 2016; Zhang et al., 2017). For instance, penguins showed positive selection in the complex I (ND4 gene) related to sea surface temperature; Australian songbirds experienced positive selection in complex I during the Pleistocene climate change (Ramos et al., 2018; Lamb et al., 2018); and high-altitude Galliforms displayed positive selection in genes of complexes I and V, adapting to increased oxygen intake (Zhou et al., 2014). Our findings support the idea that the level of positive selection on mitogenomes within taxa indicates their bioenergetic history (Garvin et al., 2015).

We identified a branch within the NWJ phylogeny under episodic positive selection likely related to the ability of NWJs to adapt their cellular metabolic needs to low elevations. The balance between heat and energy production in the OXPHOS pathway depends on the coupling efficiency of proteins involved (Portner, 2004; Wallace, 2005). A less coupled system favors heat production in cold environments, while a more coupled system prioritizes ATP production when food is scarce (Ramos et al., 2018). This suggests that positive selection along the NWJ branch may be driven by differences in proton coupling efficiency, fulfilling the energy and heat requirements specific to their environment. Future investigations should consider exploring the effects of amino acid changes using three-dimensional structures and physiochemical properties.

In conclusion, genome skimming from HTS is a valuable strategy for studying mitogenomes. Our findings in NWJ mitogenomes show a molecular shift in adaptation to low elevations where TRs and heteroplasmy could be involved. The interaction between heteroplasmic sites and TRs within the CR, possibly arising from recombination events, could potentially influence mitochondrial regulation. In contrast, these interactions in other regions may contribute to genetic polymorphism. With the advancements in HTS and LRS, we now have an exceptional opportunity to delve into mitogenomes and expand our understanding of their mechanisms and evolutionary processes.

## Author Contributions

FTG conceived the study and collected some samples. FTG and KB performed laboratory and analyzed the data. FTG, KB and SVE wrote the paper.

## Data Availability

The sequence data and complete mitogenomes have been submitted to Genbank under accession numbers [provided upon publication]. All data sets will be deposited in Dryad database at URL [available upon acceptance].

## Statements and Declarations

## Acknowledgments

We thank Emily Wheeler for comments on the manuscript. We also thank the Field Museum of Natural History (FMNH), the Biodiversity Institute at University of Kansas and the Museum of Comparative Zoology at Harvard University for providing samples.

## Supplementary Information

### Figures

**Fig.S1.** Descriptive statistics of reads sequenced and assemblies for each NWJ sample across two categories of sequencing coverage. a) Number of final assembled reads per sample, categorized by sequencing coverage (∼25X and ∼5X). The gray area represents a zoomed-in view; b) Mean organelle coverage of the final mitogenomes assembled using MITObim, AWA, and NOVOPlasty. The gray area represents a zoomed-in view. Values for MITObim and AWA were averaged across the four partitions from each individual

**Fig.S2.** Descriptive statistics of mitogenome assemblies for each NWJ sample across two categories of sequencing coverage. a) Final mitogenome length obtained by each assembler program per sample; b) Percentage of mitochondrial DNA among the total reads, calculated for MITObim, AWA, and NOVOPlasty. For MITObim and AWA, the percentage was averaged across the four partitions from each individual.

**Fig.S3.** Comparison of base composition in NWJ mitogenomes to highlight species differences and intraspecific similarities. The scatterplot displays the percentage of AT versus GC content in the mitogenomes of nine NWJ species (17 samples), with the Eurasian magpie as the outgroup. The plot illustrates the variations in base composition among the species, emphasizing both the differences between species and the similarities within species.

**Fig.S4.** Phylogenetic trees inferred from MrBayes for all mitogenomes datasets; PCGs, whole mitogenome, the control region (CR) and mitoribosomes (rRNAs); with all samples and just one sample per species of NWJs. The support for each clade is indicated at each node, with posterior probabilities provided for all clades.

**Fig.S5.** Overall genetic differences and phylogenetic relationships based on NWJ mitogenomes.

a) Pairwise genetic differences across the entire mitogenomes, illustrating the overall genetic variation between species and the reliability of mitogenomes; b) Phylogeny used for downstream analyses, constructed using PCGs and generated with MrBayes. The phylogeny includes only one sample per species, providing an overview of the species’ evolutionary relationships.

**Fig.S6.** Comparison of heteroplasmy levels in NWJ mitogenomes. The scatterplot illustrates the total number of heteroplasmic sites in all NWJs, with an increasing amount of memory/reads utilized. A plateau is observed when approximately 70% of reads are used. The Unicolored jay (*Aphelocoma unicolor*) exhibits the highest heteroplasmy levels, with 21 sites. A zoomed-in plot on the right focuses on genomes with low heteroplasmy levels and includes the three low-coverage genomes. These three low-coverage genomes show a consistent number of heteroplasmic sites from the initial run using the total reads.

**Fig.S7.** Examples of heteroplasmic sites in NWJ alignments (in purpule squares). a) One heteroplasmic site located at the beginning of the control region (CR), appearing in the same position or within a 1bp distance in *Cyanocorax* lineage, Brown jay (*Psilorhinus morio*), and Unicolored jay;

b) Heteroplasmic site identified within the NAD2 gene of Steller’s jay (*Cyanocitta stelleri*), Unicolored jay, and Azure-hooded jay (*Cyanolyca cucullata*), showing a shift in location by the red arrows.

**Fig.S8.** More examples of heteroplasmic sites in NWJ alignments (in purpule squares and pointed by the red arrows). a) Heteroplasmic site located in the NAD5 gene, approximately 90 bases apart, observed between the Bushy-crested jay (*Cyanocorax melanocyaneus*) and the Azure-hooded jay. The presence of a tandem repeat is indicated by the blue arrow; b) Heteroplasmic site identified in the COX1 gene, with a distance of 57 bases, observed between the Unicolored jay and the Azure-hooded jay; c) Heteroplasmic site found in the tRNA-L, with a distance of 20 bases, observed between the Green jay (*Cyanocorax yncas*) and the Bushy-crested jay.

**Fig.S9.** Overall TR composition in NWJ mitogenomes with the Eurasian magpie as the outgroup. a) TRs mean entropy score with error bars in all assembled mitogenomes; b) Total number of TR sites in the NWJ mitogenomes. The nucleotide percentages of TRs within the mitogenomes are indicated above the bars; c) Heatmap illustrating comparative analyses of TRs, including the total number, mean length (bp), and mean copy number (tandem unit). The heatmap reveals a pattern where NWJ species with lower numbers of TRs tend to exhibit longer TRs.

**Fig.S10.** Overall pattern of negative selection across all mitochondrial PCGs in NWJs. a) Ka/Ks ratio represented by the omega (ω) value in all PCGs, with ATP8 exhibiting the highest ω value and COX1 displaying the lowest ω value; b) Percentage of sites under negative selection in each PCG. The red dashed line indicates that 60% of the sites in all genes are under negative selection.

### Tables

**Table S1.** Tissue samples used for mitogenome assembly and analyses in 17 individuals of nine NWJ species, representing seven genera. The table presents information including the voucher, sample IDs, age, sex, location, geographic position, elevation, tissue type, and preservation conditions for each individual.

**Table S2.** Sequencing information per sample, including the total number of reads obtained, final reads assembled into mitogenomes, and the percentage of mitochondrial DNA (mtDNA) for each whole-genome library. The table also indicates the partition (if available) used by MITObim/AWA.

**Table S3.** Mitogenome lengths, average organelle coverage, and alignment scores (AWA) obtained using three assembly procedures: MITObim, AWA, and NOVOPlasty. The table presents these values for each sample and the corresponding partition sub-samples used in MITObim and AWA.

**Table S4.** Phylogenetic generalized least squares (PGLS) regression models for genome length and AT/GC base composition, as well as TR length in tRNA-P, with elevation. The regression models were conducted under the Brownian model (BM) and Ornstein-Uhlenbeck model (OU). Bold font indicates significant regressions.

**Table S5.** Results from the CodeML site model implemented in PAML, comparing model M0 (null model, rejected) with alternative models M1a, M3, and M8. The table includes the number of parameters estimated for each model (Np) and the log-likelihood ratio test p-values (P(ΔLRT)) obtained from the model comparisons. Null models are indicated in brackets.

**Table S6.** Statistically significant sites under positive selection across 5 PCGs of the NWJ mitogenomes, identified using three methods. The table displays the amino acids present at each site for each species. Significant support for each method is highlighted in red, while marginally significant support is shown in black and bold. “pp” represents the posterior probabilities.

