## Supplemental Tables S1-S6 for "Heteroplasmy and tandem repeats reveal adaptation to elevation in the New World Jays (Aves: Corvidae)"

**Table S1.** Tissue samples used for mitogenome assembly and analyses in 17 individuals of nine NWJ species, representing seven genera. The table presents information including the voucher, sample IDs, age, sex, location, geographic position, elevation, tissue type, and preservation conditions for each individual.

| Catalog_number | Species | Common_name | ID-analysis | Barcode | Field_id | Age | Sex |
|---|---|---|---|---|---|---|---|
| 343591 | *Aphelocoma unicolor* | Unicolored jay | Aph_unico | NA | MXJ-450 | NA | male |
| 109294 | *Cyanocorax melanocyaneus* | Bushy-crested jay | Cya_melan | NA | SLA-231 | NA | female |
| 334863 | *Cyanocitta stelleri* | Steller’s jay | Cya_stell | NA | ORJ-065 | NA | male |
| CRYO-8794 | *Cyanocorax yncas* | Green jay 1 | Cya_yncas_1 | CCAACA | FTG-18-301 | juvenile | male |
| ORN-365273 | *Cyanocorax yncas* | Green jay 2 | Cya_yncas_2 | CGGAAT | FTG-18-302 | juvenile | male |
| CRYO-8795 | *Cyanocorax yncas* | Green jay 3 | Cya_yncas_3 | CTCAGA | FTG-18-304 | juvenile | female |
| CRYO-8783 | *Cyanocorax yucatanicus* | Yucatan jay 1 | Cya_yucat_1 | CACGAT | FTG-18-277 | adult | male |
| ORN-365268 | *Cyanocorax yucatanicus* | Yucatan jay 2 | Cya_yucat_2 | CACTCA | FTG-18-287 | subadult | male |
| ORN-365269 | *Cyanocorax yucatanicus* | Yucatan jay 3 | Cya_yucat_3 | CAGGCG | FTG-18-292 | juvenile | female |
| ORN-365270 | *Cyanocorax yucatanicus* | Yucatan jay 4 | Cya_yucat_4 | CATGGC | FTG-18-295 | adult | male |
| CRYO-8792 | *Cyanocorax yucatanicus* | Yucatan jay 5 | Cya_yucat_5 | CATTTT | FTG-18-296 | juvenile | male |
| ORN-365274 | *Psilorhinus morio* | Brown jay 1 | Psi_morio_1 | CTATAC | FTG-18-303 | subadult | female |
| ORN-365276 | *Psilorhinus morio* | Brown jay 2 | Psi_morio_2 | GACGAC | FTG-18-306 | adult | female |
| CRYO-8796 | *Psilorhinus morio* | Brown jay 3 | Psi_morio_3 | TAATCG | FTG-18-309 | adult | male |
| 394011 | *Cyanolyca cucullata mitrata* | Azure-hooded jay | Cya_cucul | na | BMM-125 | unknown | male |
| 343604 | *Calocitta formosa colliei* | Black-throated magpie jay | Cal_colli | na | MXJ-338 | unknown | male |
| 393750 | *Gymnorhinus cyanocephalus* | Pinyon jay | Gym_cyano | na | MXJ-283 | unknown | male |

| Continuation of Table S1. | | | | | | | |
|---|---|---|---|---|---|---|---|
| Catalog_number | Lat | Long | Location | Elevation (m) | tissue type | storage_type | Collection |
| 343591 | 17.17 | -93.15 | Mexico, Guerrero, Sierra de Atoyac | 2200 | muscle | flash frozen | FMNH (The Field Museum of Natural History) |
| 109294 | 14.41 | -89.41 | El Salvador, El Limo | 1100 | muscle | ethanol | UK (University of Kansas) |
| 334863 | 44.15 | -122.57 | North America, USA, Oregon | 2001 | muscle | ethanol | FMNH |
| CRYO-8794 | 20.19 | -90.03 | Mexico, Campeche | 32 | muscle | RNAlater | MCZ (Museum of Comparative Zoology at Harvard University |
| ORN-365273 | 20.19 | -90.03 | Mexico, Campeche | 32 | muscle | RNAlater | MCZ |
| CRYO-8795 | 20.04 | -89.32 | Mexico, Yucatan | 164 | muscle | RNAlater | MCZ |
| CRYO-8783 | 21.02 | -89.5 | Mexico, Yucatan | 10 | muscle | RNAlater | MCZ |
| ORN-365268 | 20.52 | -89.37 | Mexico, Yucatan | 18 | muscle | RNAlater | MCZ |
| ORN-365269 | 20.01 | -89.01 | Mexico, Yucatan | 42 | muscle | RNAlater | MCZ |
| ORN-365270 | 20.01 | -89.01 | Mexico, Yucatan | 41 | muscle | RNAlater | MCZ |
| CRYO-8792 | 20.52 | -89.37 | Mexico, Yucatan | 12 | muscle | RNAlater | MCZ |
| ORN-365274 | 20.04 | -89.32 | Mexico, Yucatan | 164 | muscle | RNAlater | MCZ |
| ORN-365276 | 20.06 | -89.33 | Mexico, Yucatan | 165 | muscle | RNAlater | MCZ |
| CRYO-8796 | 20.05 | -89.31 | Mexico, Yucatan | 154 | muscle | SDS and RNAlater | MCZ |
| 394011 | 20.99 | -98.66 | Mexico, Hidalgo, Tlanchinol, 5 km E | 1539 | muscle | flash frozen | FMNH |
| 343604 | 25.48 | -107.94 | Mexico, Sinaloa | 180 | muscle | flash frozen | FMNH |
| 393750 | 31.04 | -115.47 | Mexico, Baja California Norte | 3000 | muscle | flash frozen | FMNH |

**Table S2.** Sequencing information per sample, including the total number of reads obtained, final reads assembled into mitogenomes, and the percentage of mitochondrial DNA (mtDNA) for each whole-genome library. The table also indicates the partition (if available) used by MITObim/AWA.

| Sample_partition | Sequencing platform | Mitobim/AWA input_reads | #Mitobim_used_reads | Mitobim_percentage_mtDNA | Awa_used_reads | Awa_percentage_mtDNA |
|---|---|---|---|---|---|---|
| *A. unicolor 01* | Illumina HiSeq 125 bp, pair-end | 30,006,240 | 512,834 | 1.71 | 514,258 | 1.71 |
| *A. unicolor 02* | | 30,006,240 | 512,771 | 1.71 | 514,209 | 1.71 |
| *A. unicolor 03* | | 30,006,240 | 514,279 | 1.71 | 515,220 | 1.72 |
| *A. unicolor 04* | | 30,006,240 | 513,304 | 1.71 | 515,380 | 1.72 |
| *C. colliei* | NovaSeq 150 bp, pair-end | 1,246,070 | 3,937 | 0.31 | 3,926 | 0.31 |
| *C. melanocyaneus 01* | Illumina HiSeq 125 bp, pair-end | 31,523,004 | 32,866 | 0.1 | 32,785 | 0.1 |
| *C. melanocyaneus 02* | | 31,523,004 | 33,200 | 0.11 | 33,145 | 0.11 |
| *C. melanocyaneus 03* | | 31,523,004 | 33,468 | 0.11 | 33,365 | 0.11 |
| *C. melanocyaneus 04* | | 31,523,004 | 33,404 | 0.11 | 33,342 | 0.11 |
| *C. stelleri 01* | Illumina HiSeq 125 bp, pair-end | 36,609,782 | 140,552 | 0.38 | 140,214 | 0.38 |
| *C. stelleri 02* | | 36,609,782 | 140,157 | 0.38 | 140,381 | 0.38 |
| *C. stelleri 03* | | 36,609,782 | 139,978 | 0.38 | 140,711 | 0.38 |
| *C. stelleri 04* | | 36,609,782 | 140,995 | 0.39 | 141,038 | 0.39 |
| *C. yncas CCAACA 01* | NovaSeq 150 bp, pair-end | 33,711,928 | 405,109 | 1.2 | 431,880 | 1.28 |
| *C. yncas CCAACA 02* | | 33,711,928 | 427,639 | 1.27 | 429,409 | 1.27 |
| *C. yncas CCAACA 03* | | 33,711,928 | 435,761 | 1.29 | 434,025 | 1.29 |
| *C. yncas CCAACA 04* | | 33,711,928 | 410,566 | 1.22 | 432,619 | 1.28 |
| *C. yncas CGGAAT 01* | | 45,820,238 | 846,771 | 1.85 | 850,303 | 1.86 |
| *C. yncas CGGAAT 02* | | 45,820,238 | 846,611 | 1.85 | 832,179 | 1.82 |
| *C. yncas CGGAAT 03* | | 45,820,238 | 808,492 | 1.76 | 832,562 | 1.82 |
| *C. yncas CGGAAT 04* | | 45,820,238 | 828,687 | 1.81 | 836,013 | 1.82 |
| *C. yncas CTCAGA 01* | | 47,118,132 | 350,148 | 0.74 | 356,766 | 0.76 |
| *C. yncas CTCAGA 02* | | 47,118,132 | 329,375 | 0.7 | 356,324 | 0.76 |
| *C. yncas CTCAGA 03* | | 47,118,132 | 327,726 | 0.7 | 356,442 | 0.76 |
| *C. yncas CTCAGA 04* | | 47,118,132 | 331,165 | 0.7 | 358,669 | 0.76 |
| *C. yucatanicus CACGAT 01* | NovaSeq 150 bp, pair-end | 42,193,000 | 943,249 | 2.24 | 967,787 | 2.29 |
| *C. yucatanicus CACGAT 02* | | 42,193,000 | 918,656 | 2.18 | 965,564 | 2.29 |
| *C. yucatanicus CACGAT 03* | | 42,193,000 | 898,330 | 2.13 | 964,951 | 2.29 |
| *C. yucatanicus CACGAT 04* | | 42,193,000 | 929,347 | 2.2 | 967,166 | 2.29 |

| | | | | | | |
|---|---|---|---|---|---|---|
| *C. yucatanicus CACTCA 01* | | 32,030,638 | 610,782 | 1.91 | 727,741 | 2.27 |
| *C. yucatanicus CACTCA 02* | | 32,030,638 | 619,090 | 1.93 | 705,982 | 2.2 |
| *C. yucatanicus CACTCA 03* | | 32,030,638 | 606,475 | 1.89 | 611,511 | 1.91 |
| *C. yucatanicus CACTCA 04* | | 32,030,638 | 580,660 | 1.81 | 610,811 | 1.91 |
| *C. yucatanicus CAGGCG 01* | | 28,930,764 | 598,908 | 2.07 | 595,918 | 2.06 |
| *C. yucatanicus CAGGCG 02* | | 28,930,764 | 601,983 | 2.08 | 598,031 | 2.07 |
| *C. yucatanicus CAGGCG 03* | | 28,930,764 | 571,263 | 1.97 | 596,213 | 2.06 |
| *C. yucatanicus CAGGCG 04* | | 28,930,764 | 572,480 | 1.98 | 596,239 | 2.06 |
| *C. yucatanicus CATGGC 01* | | 32,164,562 | 700,778 | 2.18 | 711,340 | 2.21 |
| *C. yucatanicus CATGGC 02* | | 32,164,562 | 674,149 | 2.1 | 713,446 | 2.22 |
| *C. yucatanicus CATGGC 03* | | 32,164,562 | 671,986 | 2.09 | 712,170 | 2.21 |
| *C. yucatanicus CATGGC 04* | | 32,164,562 | 704,418 | 2.19 | 712,042 | 2.21 |
| *C. yucatanicus CATTTT 01* | | 39,968,356 | 469,669 | 1.18 | 476,934 | 1.19 |
| *C. yucatanicus CATTTT 02* | | 39,968,356 | 450,339 | 1.13 | 477,211 | 1.19 |
| *C. yucatanicus CATTTT 03* | | 39,968,356 | 456,388 | 1.14 | 476,520 | 1.19 |
| *C. yucatanicus CATTTT 04* | | 39,968,356 | 463,605 | 1.16 | 477,461 | 1.19 |
| *C. cucullata* | NovaSeq 150 bp, pair-end | 920,616 | 1,808 | 0.19 | 1,801 | 0.19 |
| *G. cyanocephalus* | NovaSeq 150 bp, pair-end | 1,155,288 | 2,481 | 0.21 | 2,477 | 0.21 |
| *P. morio CTATAC 01* | NovaSeq 150 bp, pair-end | 37,421,166 | 740,536 | 1.98 | 835,823 | 2.23 |
| *P. morio CTATAC 02* | | 37,421,166 | 742,956 | 1.99 | 838,544 | 2.24 |
| *P. morio CTATAC 03* | | 37,421,166 | 767,960 | 2.05 | 838,443 | 2.24 |

| | | | | | |
|---|---|---|---|---|---|
| *P. morio CTATAC 04* | 37,421,166 | 767,359 | 2.05 | 834,948 | 2.23 |
| *P. morio GACGAC 01* | 36,441,115 | 413,804 | 1.14 | 425,643 | 1.17 |
| *P. morio GACGAC 02* | 36,441,115 | 414,680 | 1.14 | 425,324 | 1.17 |
| *P. morio GACGAC 03* | 36,441,115 | 412,029 | 1.13 | 428,856 | 1.18 |
| *P. morio GACGAC 04* | 36,441,115 | 425,348 | 1.17 | 425,030 | 1.17 |
| *P. morio TAATCG 01* | 38,167,274 | 103,116 | 0.27 | 102,215 | 0.27 |
| *P. morio TAATCG 02* | 38,167,274 | 97,374 | 0.26 | 101,916 | 0.27 |
| *P. morio TAATCG 03* | 38,167,274 | 111,130 | 0.29 | 102,020 | 0.27 |
| *P. morio TAATCG 04* | 38,167,274 | 101,057 | 0.26 | 102,535 | 0.27 |

**Continuation of Table S2.**

| Sample-no-partition | Novoplasty_total_reads | Novoplasty_assembled_reads | Novoplasty_percentage_mtDNA |
|---|---|---|---|
| *A. unicolor* | 120,024,960 | 250,838 | 1.64 |
| *C. colliei* | 120,024,960 | 250,838 | 1.64 |
| *C. melanocyaneus* | 120,024,960 | 250,838 | 1.64 |
| *C. stelleri* | 120,024,960 | 250,838 | 1.64 |
| *C. yncas CCAACA* | 134,847,710 | 1,432,366 | 1.18 |
| *C. yncas CGGAAT* | 134,847,710 | 1,432,366 | 1.18 |
| *C. yncas CTCAGA* | 134,847,710 | 1,432,366 | 1.18 |
| *C. yucatanicus CACGAT* | 134,847,710 | 1,432,366 | 1.18 |
| *C. yucatanicus CACTCA* | 128,122,552 | 2,057,552 | 1.76 |
| *C. yucatanicus CAGGCG* | 128,122,552 | 2,057,552 | 1.76 |
| *C. yucatanicus CATGGC* | 128,122,552 | 2,057,552 | 1.76 |
| *C. yucatanicus CATTTT* | 128,122,552 | 2,057,552 | 1.76 |
| *C. cucullata* | 920,616 | 2,340 | 0.27 |
| *G. cyanocephalus* | 1,155,288 | 3,040 | 0.29 |
| *P. morio CTATAC* | 149,684,660 | 2,736,020 | 2.03 |
| *P. morio GACGAC* | 149,684,660 | 2,736,020 | 2.03 |
| *P. morio TAATCG* | 149,684,660 | 2,736,020 | 2.03 |

**Table S3.** Mitogenome lengths, average organelle coverage, and alignment scores (AWA) obtained using three assembly procedures: MITObim, AWA, and NOVOPlasty. The table presents these values for each sample and the corresponding partition sub-samples used in MITObim and AWA.

| Sample_partition | Mitobim_avg_coverage | Mitobim_mitogenome_length | Awa_avg_coverage | Awa_mitogenome_length | AWA_avg_alignment_score |
|---|---|---|---|---|---|

| | | | | | |
|---|---|---|---|---|---|
| *A. unicolor 01* | 3,703.54 | 17,421 | 2,495.44 | 16,910 | -0.79 |
| *A. unicolor 02* | 3,707.67 | 17,402 | 2,576.14 | 16,903 | -1.18 |
| *A. unicolor 03* | 3,696.21 | 17,527 | 2,526.65 | 16,900 | -0.81 |
| *A. unicolor 04* | 3,556.67 | 18,313 | 129.21 | 17,304 | -3.776* |
| *C. colliei* | 37.63 | 17,164 | 37.63 | 16,888 | -0.97 |
| *C. melanocyaneus 01* | 242.68 | 17,277 | 248.59 | 16,911 | -0.48 |
| *C. melanocyaneus 02* | 246.9 | 17,158 | 244.62 | 16,911 | -0.29 |
| *C. melanocyaneus 03* | 244.39 | 17,424 | 235.41 | 16,911 | -0.46 |
| *C. melanocyaneus 04* | 248.24 | 17,164 | 234.94 | 16,911 | -0.41 |
| *C. stelleri 01* | 1,014.40 | 17,424 | 688.86 | 16,906 | -8.553* |
| *C. stelleri 02* | 1,016.78 | 17,340 | 191.41 | 16,904 | -2.43 |
| *C. stelleri 03* | 1,004.54 | 17,604 | 858.1 | 17,012 | -0.81 |
| *C. stelleri 04* | 1,021.87 | 17,348 | 710.8 | 16,905 | -0.71 |
| *C. yncas CCAACA 01* | 3,453.21 | 17,670 | 3,407.14 | 16,914 | -0.44 |
| *C. yncas CCAACA 02* | 3,535.85 | 18,177 | 3,427.67 | 16,932 | -0.48 |
| *C. yncas CCAACA 03* | 3,683.13 | 17,598 | 3,399.18 | 17,001 | -0.48 |
| *C. yncas CCAACA 04* | 3,477.94 | 17,808 | 3,542.39 | 16,954 | -0.56 |
| *C. yncas CGGAAT 01* | 6,569.80 | 19,302 | 17 | 18,030 | -5.705* |
| *C. yncas CGGAAT 02* | 6,711.82 | 19,071 | 6,000.99 | 16,919 | -0.46 |
| *C. yncas CGGAAT 03* | 6,636.72 | 18,336 | 854.12 | 16,928 | -2.47 |
| *C. yncas CGGAAT 04* | 6,775.32 | 18,433 | 5,590.58 | 17,031 | -6.140* |
| *C. yncas CTCAGA 01* | 2,841.71 | 18,546 | 3,496.63 | 16,936 | -0.6 |
| *C. yncas CTCAGA 02* | 2,767.99 | 18,007 | 3,553.61 | 16,953 | -0.58 |
| *C. yncas CTCAGA 03* | 2,718.21 | 18,150 | 3,313.14 | 16,922 | -0.66 |
| *C. yncas CTCAGA 04* | 2,799.52 | 17,813 | 2,920.76 | 16,961 | -0.64 |
| *C. yucatanicus CACGAT 01* | 7,692.51 | 18,478 | 7,379.78 | 17,049 | -0.53 |
| *C. yucatanicus CACGAT 02* | 7,744.26 | 17,931 | 7,575.87 | 16,948 | -0.6 |
| *C. yucatanicus CACGAT 03* | 7,480.97 | 18,053 | 7,554.62 | 16,979 | -0.57 |
| *C. yucatanicus CACGAT 04* | 7,402.00 | 19,001 | 7,690.16 | 16,942 | -0.57 |
| *C. yucatanicus CACTCA 01* | 4,826.27 | 19,108 | 112.16 | 18,578 | -0.69 |
| *C. yucatanicus CACTCA 02* | 4,841.28 | 19,385 | 13,096.02 | 17,911 | -5.330* |
| *C. yucatanicus CACTCA 03* | 5,049.47 | 18,030 | 4,841.86 | 17,003 | -0.55 |
| *C. yucatanicus CACTCA 04* | 4,891.62 | 17,872 | 4,814.70 | 16,921 | -0.6 |
| *C. yucatanicus CAGGCG 01* | 4,948.92 | 18,084 | 4,668.43 | 16,926 | -0.57 |
| *C. yucatanicus CAGGCG 02* | 4,662.70 | 19,362 | 4,329.09 | 17,133 | -4.497* |
| *C. yucatanicus CAGGCG 03* | 4,783.54 | 18,083 | 4,580.42 | 17,000 | -0.52 |
| *C. yucatanicus CAGGCG 04* | 4,751.80 | 18,191 | 4,567.57 | 17,046 | -0.55 |
| *C. yucatanicus CATGGC 01* | 5,850.53 | 17,965 | 6,018.15 | 16,924 | -0.59 |
| *C. yucatanicus CATGGC 02* | 5,411.90 | 18,986 | 1,813.37 | 17,185 | -5.040* |

| | | | | | |
|---|---|---|---|---|---|
| *C. yucatanicus CATGGC 03* | 5,566.95 | 18,459 | 4,888.95 | 17,076 | -7.777* |
| *C. yucatanicus CATGGC 04* | 5,743.29 | 18,261 | 6,081.71 | 17,088 | -0.59 |
| *C. yucatanicus CATTTT 01* | 3,787.20 | 18,806 | 80.8 | 17,531 | -1.113* |
| *C. yucatanicus CATTTT 02* | 3,840.51 | 17,691 | 3,690.75 | 16,990 | -0.54 |
| *C. yucatanicus CATTTT 03* | 3,836.33 | 18,037 | 3,761.24 | 16,938 | -0.61 |
| *C. yucatanicus CATTTT 04* | 3,860.33 | 18,102 | 3,576.68 | 17,060 | -0.58 |
| *C. cucullata* | 18.72 | 17,329 | 18.45 | 16,886 | -0.34 |
| *G. cyanocephalus* | 24.74 | 17,209 | 21.71 | 16,897 | -0.35 |
| *P. morio CTATAC 01* | 6,341.28 | 17,671 | 7,183.06 | 16,890 | -0.84 |
| *P. morio CTATAC 02* | 6,256.35 | 17,938 | 3,203.43 | 17,047 | -2.81 |
| *P. morio CTATAC 03* | 6,321.23 | 18,267 | 7,510.90 | 16,896 | -0.91 |
| *P. morio CTATAC 04* | 6,374.11 | 18,089 | 7,041.58 | 16,892 | -0.71 |
| *P. morio GACGAC 01* | 3,508.75 | 17,760 | 517.3 | 16,919 | -3.255* |
| *P. morio GACGAC 02* | 3,379.37 | 18,795 | 3,233.36 | 17,105 | -0.56 |
| *P. morio GACGAC 03* | 3,459.14 | 17,885 | 3,521.66 | 16,886 | -0.48 |
| *P. morio GACGAC 04* | 3,524.67 | 18,123 | 3,301.41 | 16,917 | -4.217* |
| *P. morio TAATCG 01* | 864.24 | 18,020 | 1,512.92 | 16,895 | -0.77 |
| *P. morio TAATCG 02* | 826.89 | 17,976 | 1,389.52 | 16,891 | -0.79 |
| *P. morio TAATCG 03* | 926.42 | 17,917 | 1,449.52 | 16,889 | -0.74 |
| *P. morio TAATCG 04* | 850.29 | 17899 | 1,421.52 | 16,888 | -0.84 |

**Continuation of Table S3.**

| Sample-no-partition | Novoplasty_avg_coverage | Novoplasty_mitogenome_length |
|---|---|---|
| *A. unicolor* | 17,574 | 16,899 |
| *C. colliei* | 48 | 16,888 |
| *C. melanocyaneus* | 1145 | 16,911 |
| *C. stelleri* | 4891 | 16,904 |
| *C. yncas CCAACA* | 14,252 | 16,913 |
| *C. yncas CGGAAT* | 25,014 | 16,914 |
| *C. yncas CTCAGA* | 10091 | 16,914 |
| *C. yucatanicus CACGAT* | 31,148 | 16,916 |
| *C. yucatanicus CACTCA* | 20,213 | 16,916 |
| *C. yucatanicus CAGGCG* | 19,833 | 16,916 |
| *C. yucatanicus CATGGC* | 23,611 | 16,915 |
| *C. yucatanicus CATTTT* | 15,851 | 16,916 |
| *C. cucullata* | 22 | 16,886 |
| *G. cyanocephalus* | 30 | 16,897 |
| *P. morio CTATAC* | 27,274 | 16,884 |
| *P. morio GACGAC* | 14,199 | 16,884 |
| *P. morio TAATCG* | 3366 | 16,883 |

**Table S4.** Phylogenetic generalized least squares (PGLS) regression models for genome length and AT/GC base composition, as well as TR length in tRNA-P, with elevation. The regression models were conducted under the Brownian model (BM) and Ornstein-Uhlenbeck model (OU). Bold font indicates significant regressions.

| Variables | Level of analysis | Model | Intercept | Slope | Corr. Coef | SE | AIC | t | Residuals Med. | Residuals SE | P-value | Alpha |
|---|---|---|---|---|---|---|---|---|---|---|---|---|
| Genome length ~ AT | species(n= 10) | BM | 15754.2170 | 20.6910 | -0.9990 | 7.2633 | **85.3539** | 2.8487 | -0.2740 | 23.9426 | 0.0215 | NA |
| | genera (n=7) | OU | 15345.9700 | 27.8570 | -1.0000 | 7.7284 | 82.2285 | 3.6045 | **0.0980** | 9.8995 | **0.0069** | 2.5272 |
| Genome length ~ GC | species(n= 10) | BM | 17755.3250 | -19.2310 | -0.9990 | 9.5322 | **88.2438** | -2.0175 | -0.3098 | 27.6647 | 0.0784 | NA |
| | genera (n=7) | OU | 18106.0310 | -27.4490 | -1.0000 | 11.5557 | 86.5742 | -2.3754 | 0.1140 | 12.3263 | 0.0449 | 23.3634 |
| Elevation ~ TR length | species(n= 10) | BM | 2077.1950 | -1815.5340 | -0.1760 | 227.5689 | **158.8368** | -7.9780 | **-0.0950** | 943.6993 | **0.0001** | NA |
| | genera (n=7) | OU | 2086.1060 | -1866.7820 | -0.4430 | 261.3397 | 156.8093 | -7.1431 | -0.1913 | 447.5419 | **0.0001** | 5.0978 |

**Table S5.** Results from the CodeML site model implemented in PAML, comparing model M0 (null model, rejected) with alternative models M1a, M3, and M8. The table includes the number of parameters estimated for each model (Np) and the log-likelihood ratio test p-values (P(ΔLRT)) obtained from the model comparisons. Null models are indicated in brackets.

| Model | Log likelihood | Np | P (ΔLRT) | Parameters | Interpretation |
|---|---|---|---|---|---|
| M0 | -42571.26 | 19 | - | ω = 0.036 | - |
| M1a | -42136.05 | 20 | < 0.001 (M0) | p0 = 0.954, p1 = 0.045<br>ω0 = 0.018, ω1 = 1.0 | ω varies – nearly neutral |
| M2a | -42136.05 | 22 | P = 0.999 (M1a) | p0 = 0.954, p1 = 0.045, p2 = 0.0<br>ω0 = 0.018, ω1 = 1.0, ω2 = 43.93 | No positive selection |
| M3 | -41958.64 | 23 | < 0.001 (M0) | p0 = 0.867, p1 = 0.126, p2 = 0.006<br>ω0 = 0.005, ω1 = 0.242, ω2 = 1.336 | Variable selection pressure is present among sites |
| M7 | -42097.18 | 20 | - | P = 0.027, q = 0.187 | - |
| M8 | -42014.56 | 22 | < 0.001 (M7) | p0 = 0.979, p1 =0.02, ω = 1<br>p = 0.028, q = 0.211 | Positive selection |

**Table S6.** Statistically significant sites under positive selection across 5 PCGs of the NWJ mitogenomes, identified using three methods. The table displays the amino acids present at each site for each species. Significant support for each method is highlighted in red, while marginally significant support is shown in black and bold. "pp" represents the posterior probabilities.

| Gene | Position | Eurasian_magpie | Azure-hooded_jay | Black-throated_magpie_jay | Brown_jay | Green_jay | Bushy-crested_jay | Yucatan_jay | Pinyon_jay | Unicolored_jay | Steller's_jay | FUBAR_pp | MEME_p_value | PAML_pp |
|---|---|---|---|---|---|---|---|---|---|---|---|---|---|---|
| ATP6 | 183 | I | L | L | L | L | T | V | V | L | L | 0.32 | 0.00 | **0.84** |
| ATP8 | 18 | I | Y | Y | Y | F | F | L | F | L | I | 0.94 | **0.06** | **0.78** |
| COX2 | 44 | L | H | A | V | F | F | L | M | M | M | 0.41 | 0.05 | 0.97 |
| ND2 | 317 | I | V | N | T | V | T | T | L | S | I | 0.90 | **0.13** | 0.98 |
| ND3 | 20 | T | L | T | M | A | T | T | L | L | M | 0.90 | **0.09** | 0.88 |
