## Supplementary figures and images for "Heteroplasmy and tandem repeats reveal adaptation to elevation in the New World Jays (Aves: Corvidae)"

### Fig.S1.

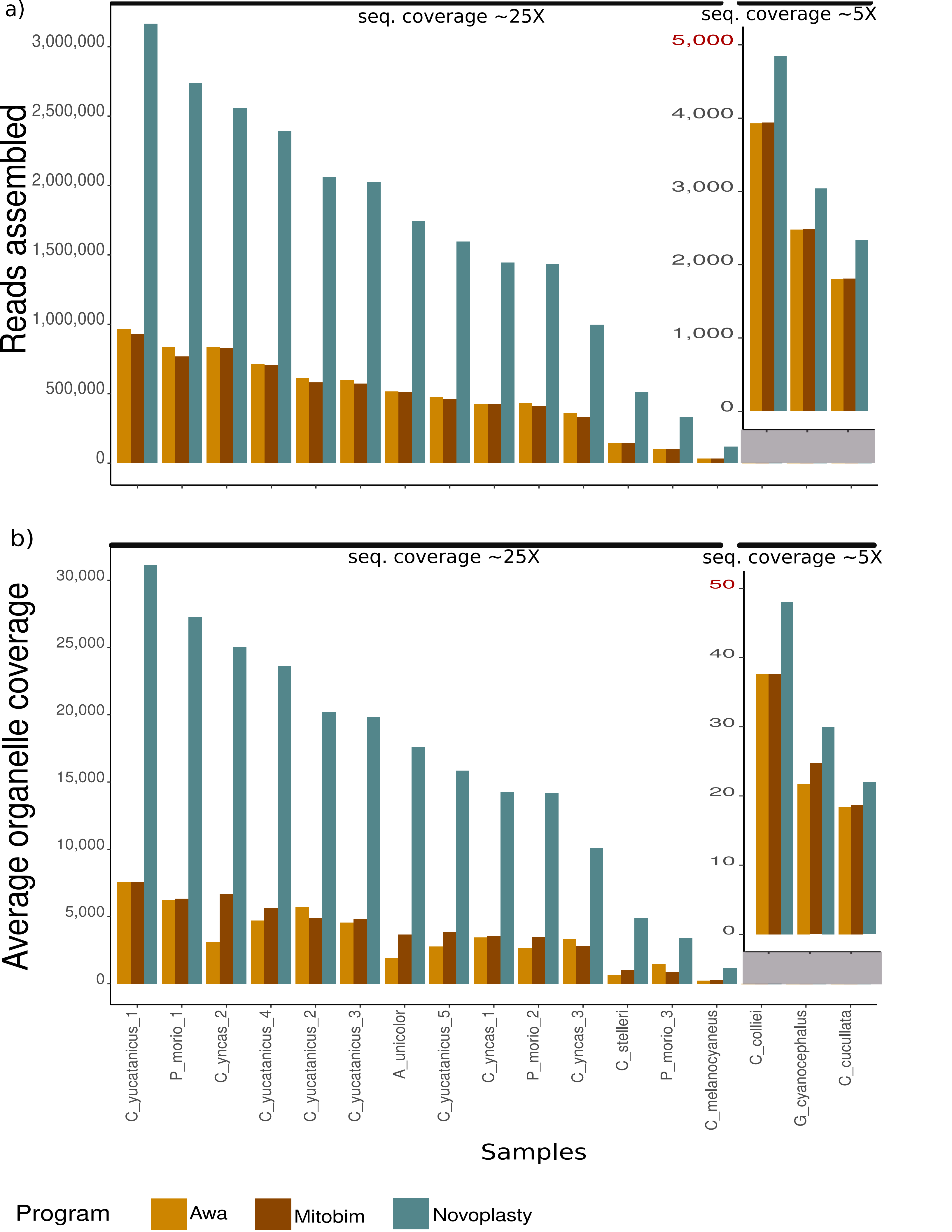

### Fig.S2.

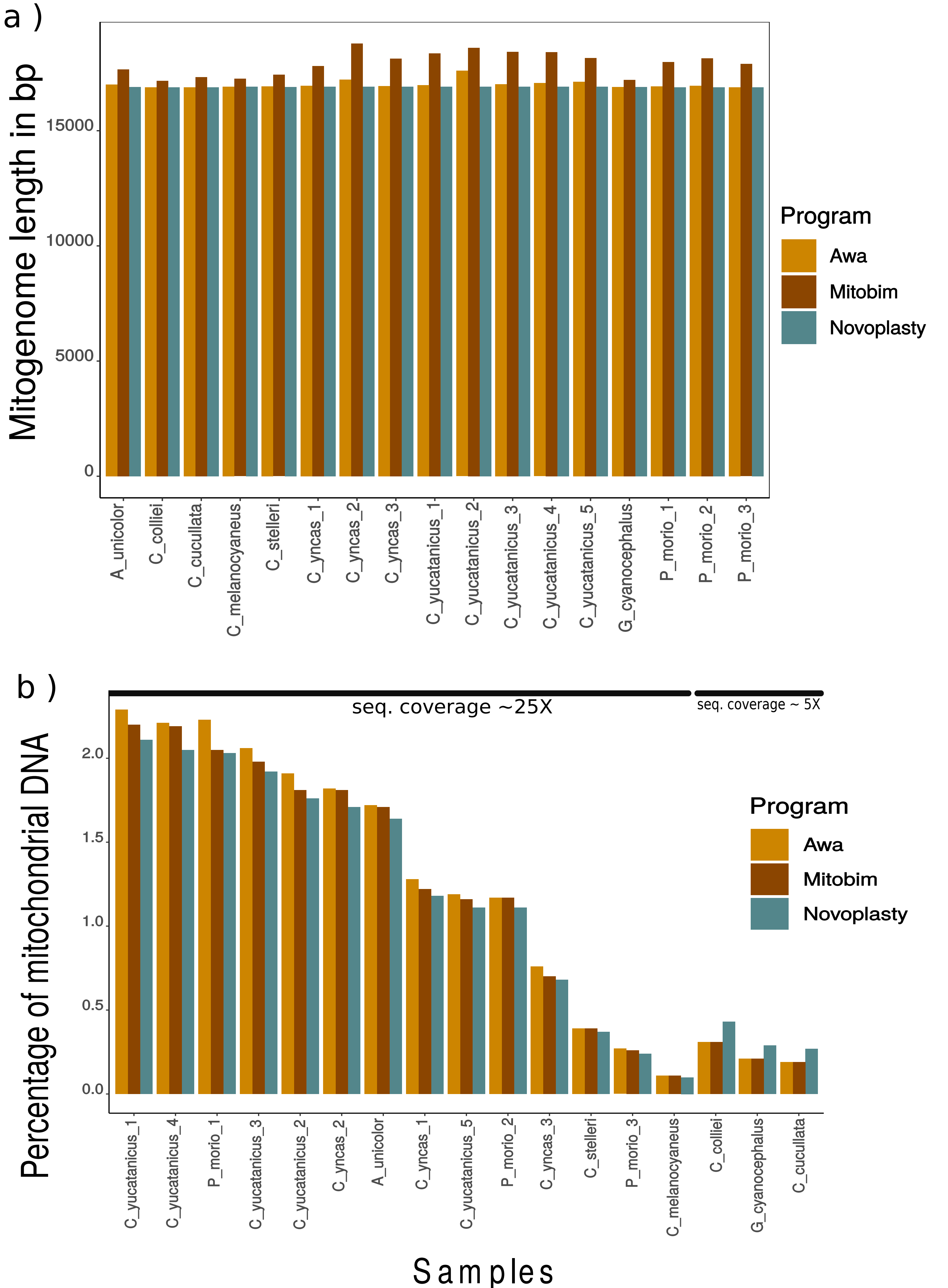

### Fig.S3.

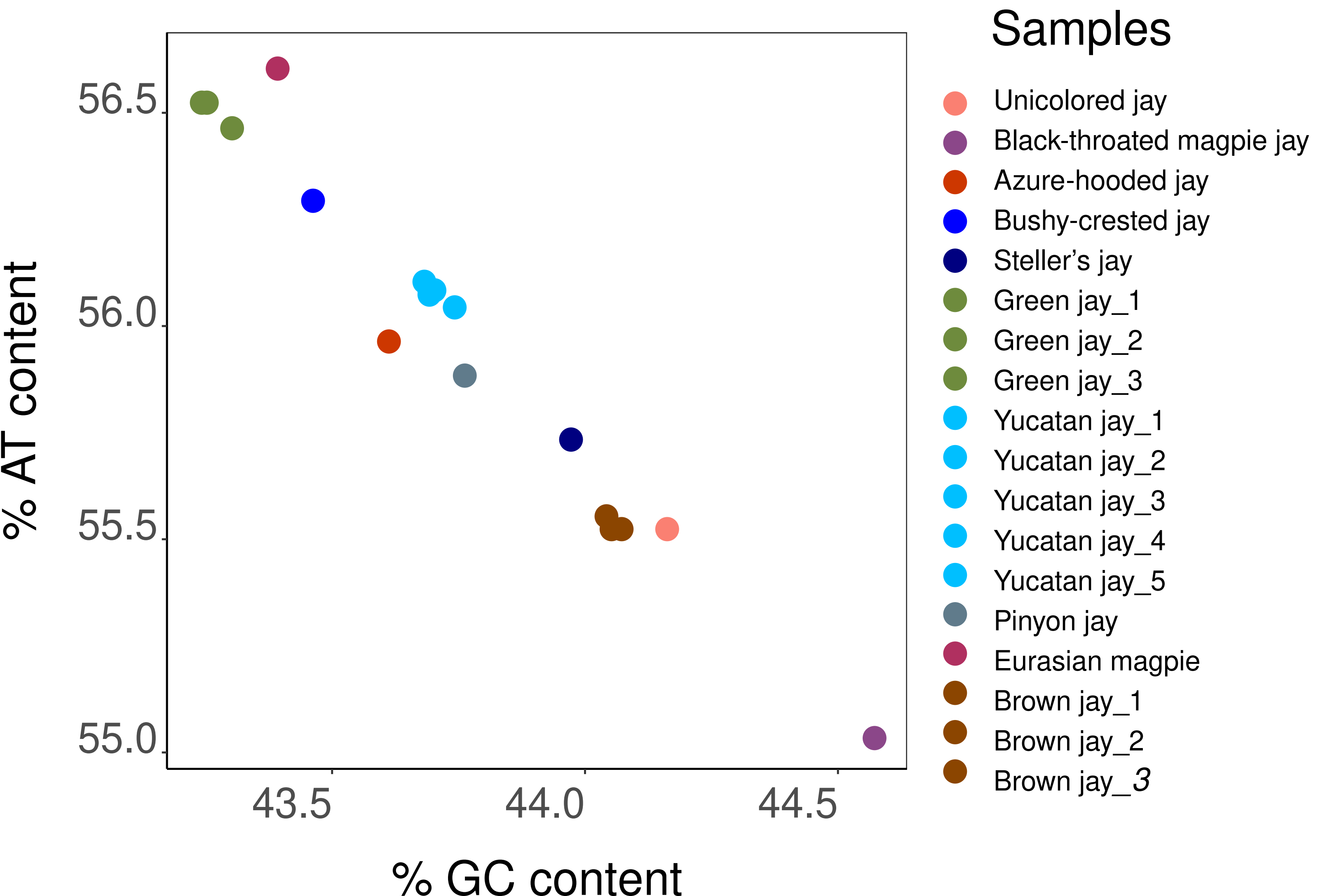

### Fig.S4.

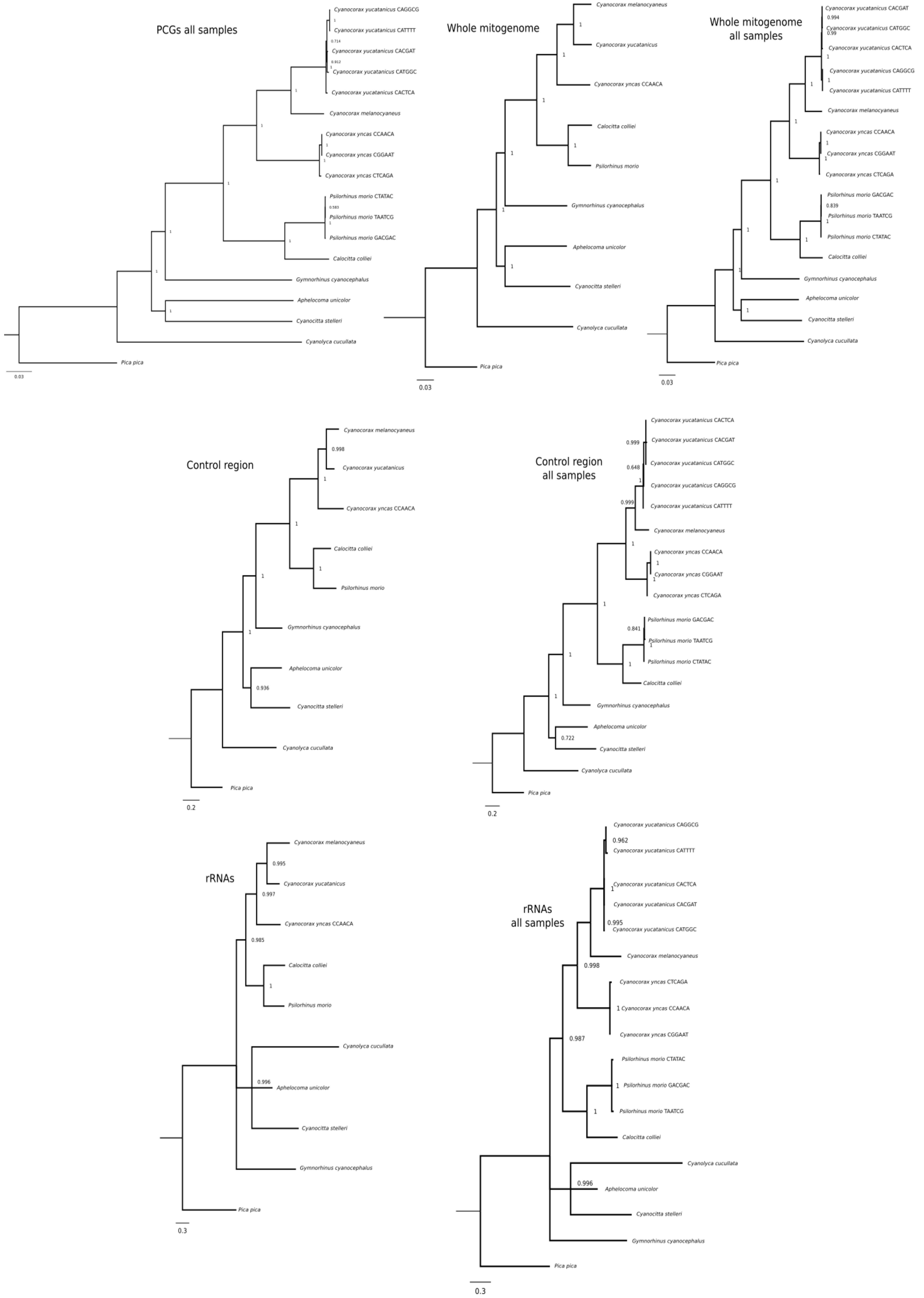

### Fig.S5.

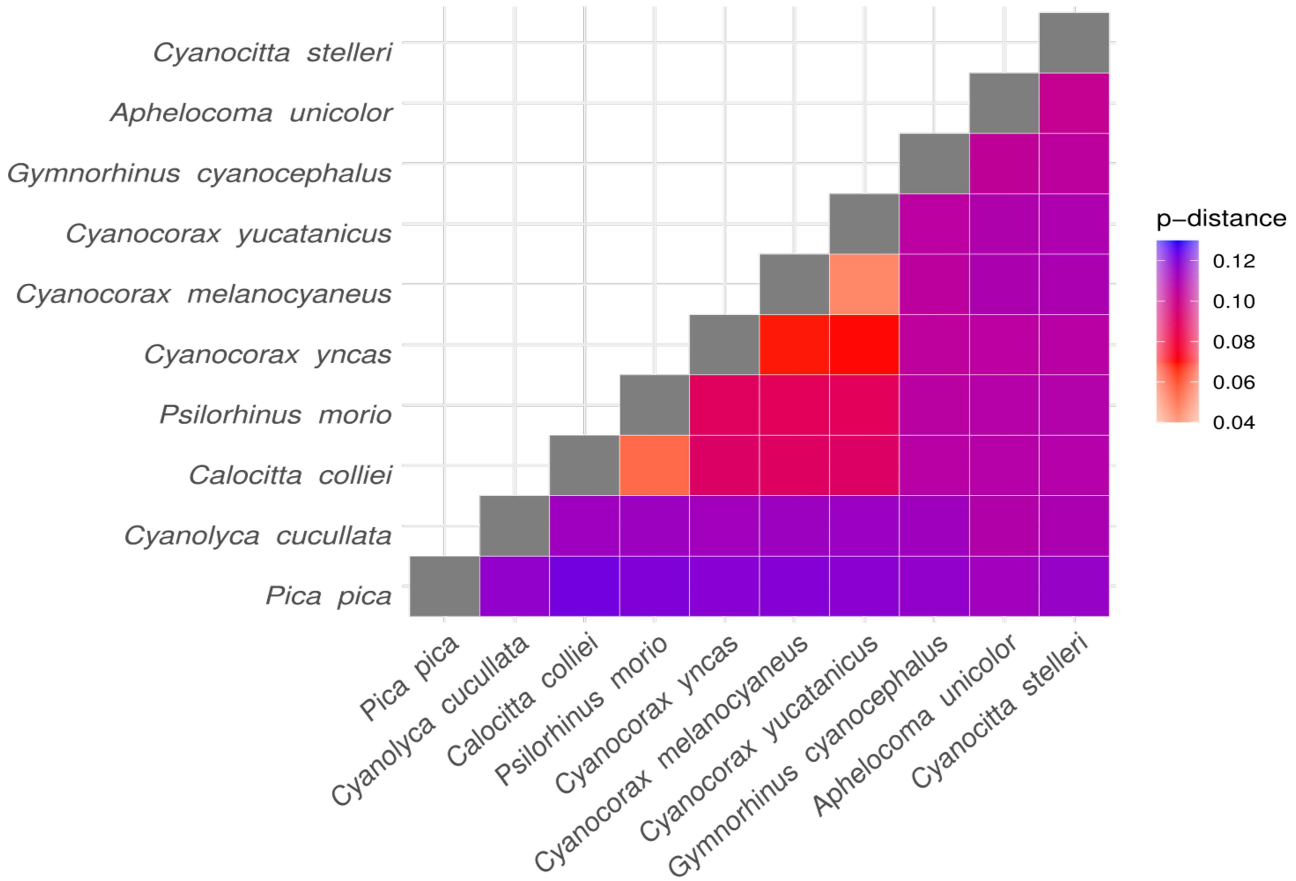

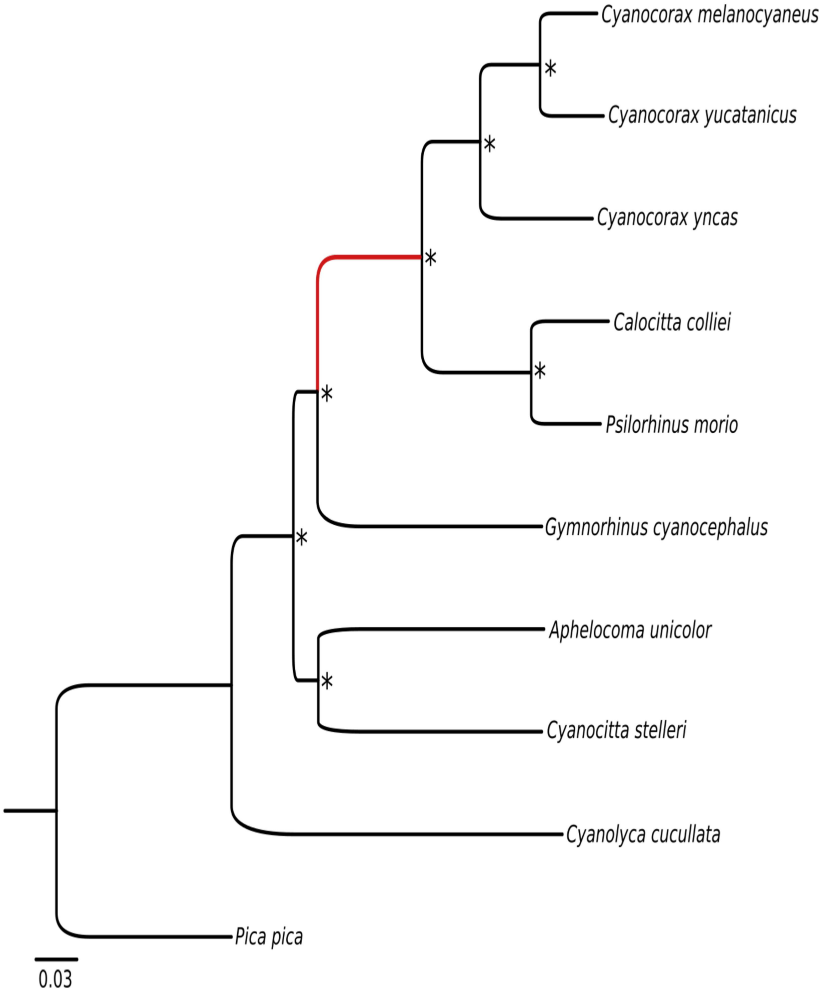

### Fig.S6.

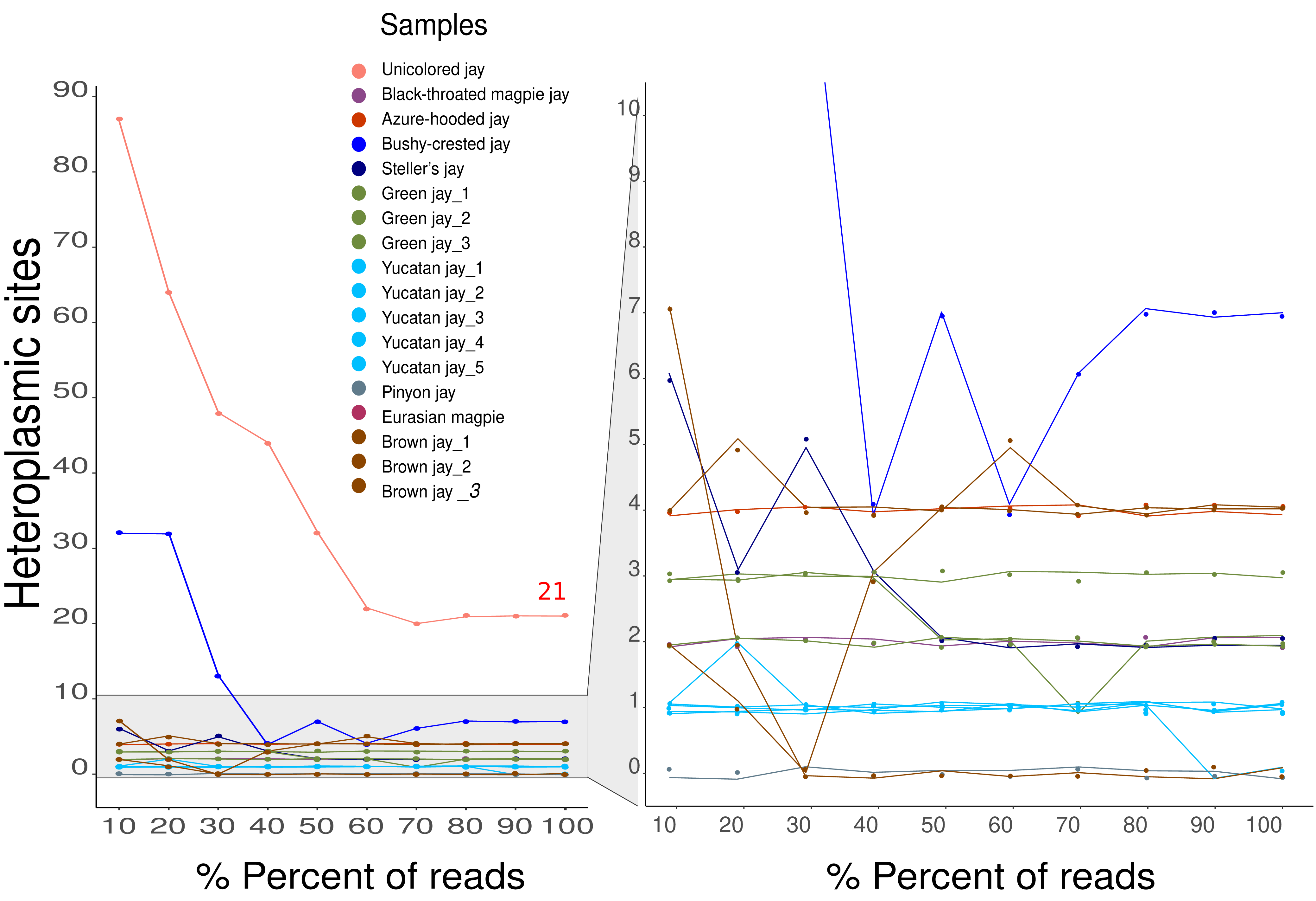

### Fig.S7.

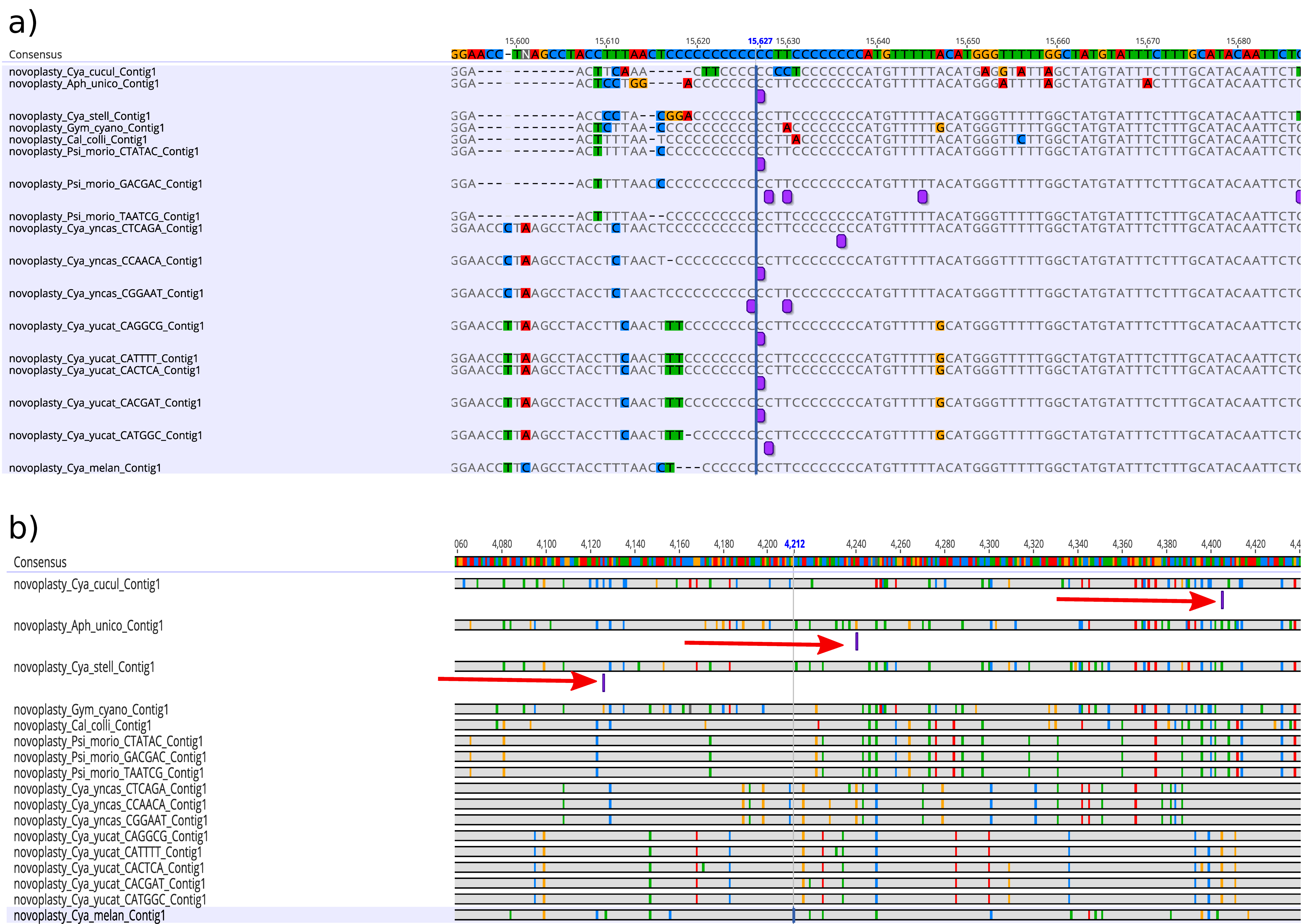

### Fig.S8.

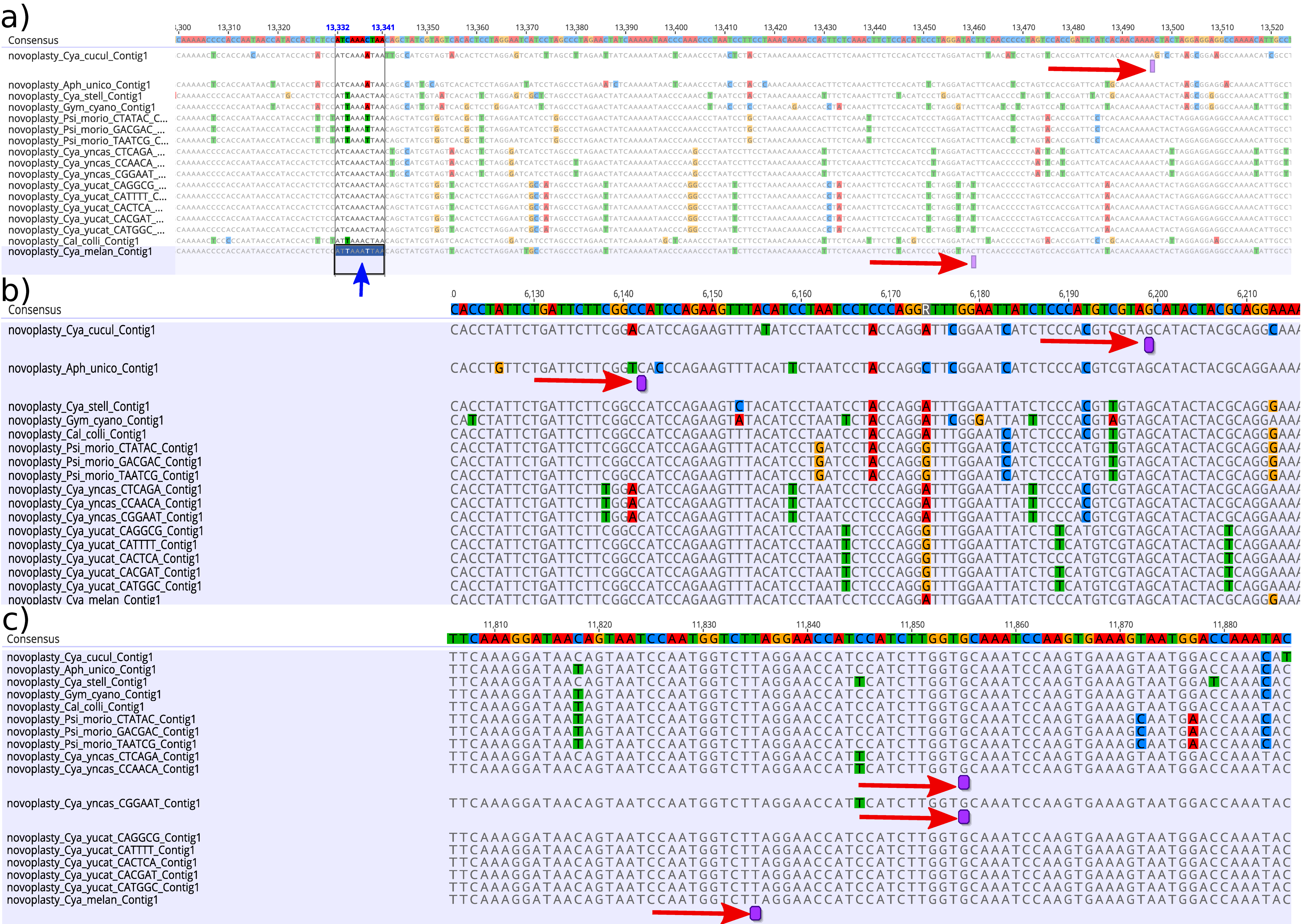

### Fig.S9.

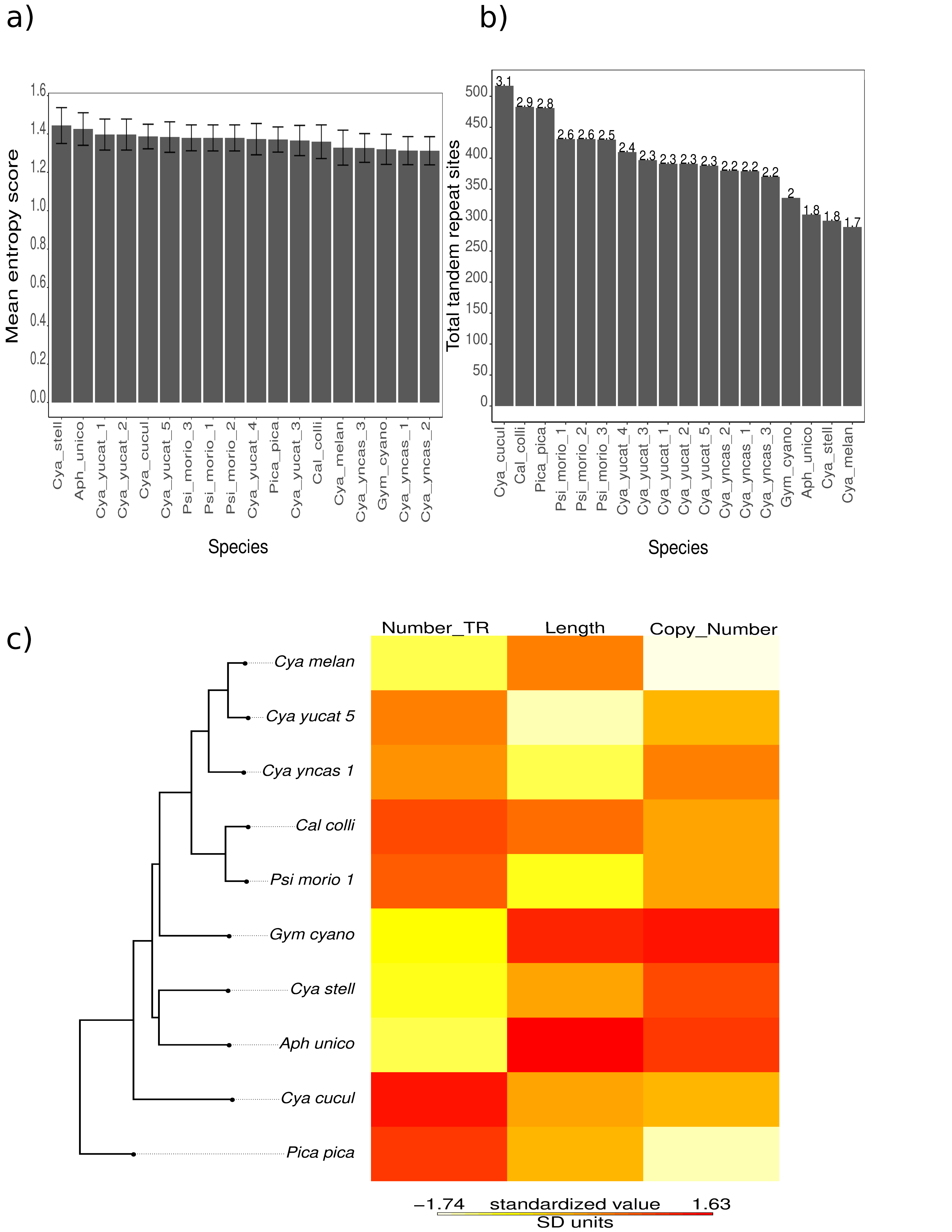

### Fig.S10.

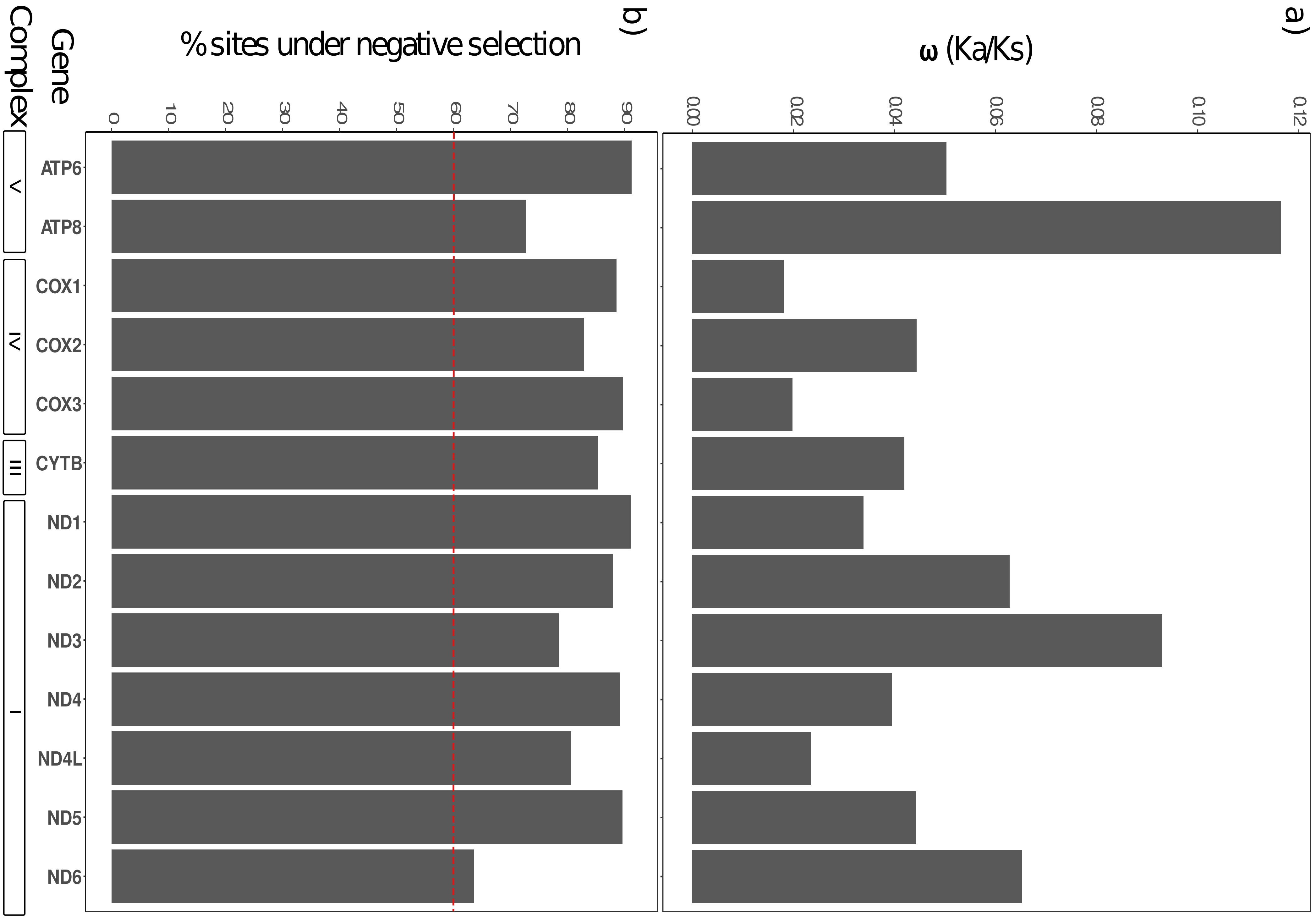
